# Biosynthesis, Structure, and Antibiotic Properties of Gelatinamin A, a Triculamin-like Lasso Peptide

**DOI:** 10.64898/2025.12.02.691826

**Authors:** Tiziana Svenningsen, Aske Merrild, Fang Wang, Frederikke D. Hansen, Toby G. Johnson, Reina Iwase, Thibault Viennet, Morten Krog Larsen, A. James Link, Thomas Tørring

## Abstract

Lasso peptides are structurally unique natural products endowed with high thermal and proteolytic stability, making them attractive as scaffolds for drug discovery. Recently, a new class of lasso peptides containing a second macrocycle, formed between a lysine sidechain and the C-terminus, was discovered, resulting in an even more compact architecture. Here, we report the first NMR structure of the class V lasso peptide, gelatinamin A. Using heterologous expression of the gelatinamin biosynthetic gene cluster (BGC) in *Bacillus subtilis*, we delineated the biosynthetic pathway through targeted gene deletions. We expressed and characterized the predicted transpeptidase, GelP, that catalyzes the formation of an isopeptide bond between Lys2 and the C-terminus and mediates the reversible conversion of gelatinamin B to gelatinamin A. In addition, we characterize GelT, an *N-*acetyltransferase that inactivates lasso peptide antimicrobial activity by acetylating a key lysine residue. Furthermore, we demonstrate that gelatinamin is highly potent against several important pathogens and that the activity is strongly bicarbonate-dependent. Finally, we propose a complete biosynthetic pathway for gelatinamin. The structural insight of gelatinamin A and the functional characterization of GelP provide the foundation for future discovery of class V lasso peptides and for engineering transpeptidases to modify other lasso peptide scaffolds.

## Introduction

Ribosomally synthesized and post-translationally modified peptides (RiPPs) constitute an intriguing class of natural products distinguished by their structural complexity, biosynthetic logic, and diverse biological functions. RiPP biosynthesis is typically governed by compact biosynthetic gene clusters (BGCs) that encode a precursor peptide along with tailoring enzymes, transport proteins, and occasionally self-resistance elements. ^[1–5]^

In recent years, genome mining combined with advances in bioinformatics have greatly expanded our knowledge of RiPPs, particularly lasso peptides.^[6–11]^ These are structurally unique, defined by their threaded lariat structure and remarkable stability. Traditionally, lasso peptides have been described as undergoing relatively simple chemical maturation, involving leader peptide cleavage and macrolactam ring formation, while exhibiting the interlocked [1]rotaxane architecture.^[12–14]^ However, an increasing number of lasso peptides have now been discovered with unusual and diverse post-translational modifications, including disulfide bond formation,^[15]^ phosphorylation,^[16]^ methylation,^[17]^ acetylation,^[18,19]^ hydroxylation,^[20]^ citrullination,^[8]^ aspartimidylation,^[21]^ and several others.^[13,22–25]^ These modifications not only enhance structural diversity and functional complexity but also challenge our current understanding of lasso peptide biosynthesis and their biological roles. The continued discovery of such modified lasso peptides underscores a broader and more intricate landscape of directed peptide modifications than previously appreciated.

Gelatinamins are novel triculamin-like lasso peptides first reported in our previous study, where it was shown to exhibit bioactivity against *Mycobacteria* strains.^[19]^ Gelatinamin A has the class defining lasso modification of an N-terminal to Asp8 isopeptide bond. Furthermore, it features two distinctive post-translational modifications: *N-*acetylation and a second macrocyclization between the carboxylate of the C-terminus and the *ε*-amino group of a lysine in the lasso ring, accompanied by the loss of glycine. These modifications are predicted to be catalyzed by an *N-*acetyltransferase (GelT) and a clostripain-like peptidase (GelP). Notably, this unusual cross-linking was also found in lariocidin B and proposed to be a class-defining hallmark of class V lasso peptides.^[26]^ Gelatinamin A and lariocidin B are the only biologically characterized peptides with this unique figure-of-eight interlocked architecture,^[27]^ expanding the structural diversity of naturally occurring mechanically interlocked peptides.

In this study, we investigate the biosynthetic pathway of gelatinamin by identifying and characterizing the minimal gene cluster required for its production. Using *Bacillus subtilis* SCK6 as a heterologous host,^[28]^ we reconstituted the biosynthetic gene cluster and performed systematic gene deletions to determine which genes are necessary and sufficient for gelatinamin formation. In addition, we used the native producer *B. gelatini* to produce gelatinamins and purified the major gelatinamin variant, gelatinamin A, and determined its three-dimensional structure using nuclear magnetic resonance (NMR) spectroscopy. The purified gelatinamin A was assessed for antimicrobial activity against *Mycobacteria*, several ESKAPE pathogens (*Staphylococcus aureus, Klebsiella pneumonia, Acinetobacter baumannii*, and *Pseudomonas aeruginosa*), and attenuated *E. coli* strains in the presence and absence of physiological concentrations of bicarbonate. The observed broad-spectrum antimicrobial activity displayed a pronounced bicarbonate-dependent enhancement with minimum inhibitory concentrations (MICs) <1 µg/mL in several cases. Finally, we reconstituted both the enzymatic acetylation and the unusual macrocyclization *in vitro* using purified GelT and GelP, enabling us to propose a complete biosynthetic pathway for gelatinamin.

## Results and Discussion

### The *gel* BGC is Necessary and Sufficient for Gelatinamin Biosynthesis and Export in *B. subtilis*

To identify the genes required for gelatinamin biosynthesis, we amplified the gelatinamin gene cluster (*gel*) from *Brevibacillus gelatini* PDF4^[29]^ genomic DNA using PCR (Figure 1a). The cluster is predicted to include a precursor (*gelA*), a standalone RiPP recognition element (RRE, *gelB1*), a peptidase (*gelB2*), a macrocyclase (*gelC*), two ABC transporters (*gelD1* and *gelD2*), a putative clostripain-like peptidase (*gelP*), and an *N-*acetyltransferase (*gelT*) as shown in Figure 1a. The amplified fragment was cloned into a pRB373-based plasmid^[30]^ using In-Fusion cloning. After purification and sequence verification of the plasmid via whole-plasmid sequencing, it was introduced into *B. subtilis* SCK6 through natural transformation, facilitated by xylose-induced activation of the ComK gene (Figure 1b).^[31]^ Growth optimization showed that TB culture medium outperformed LB (Figure S1), so all subsequent experiments used TB. This approach allowed us to test if the *gel* BGC was sufficient for heterologous production in *B. subtilis* and which genes are necessary.

**Figure 1.**
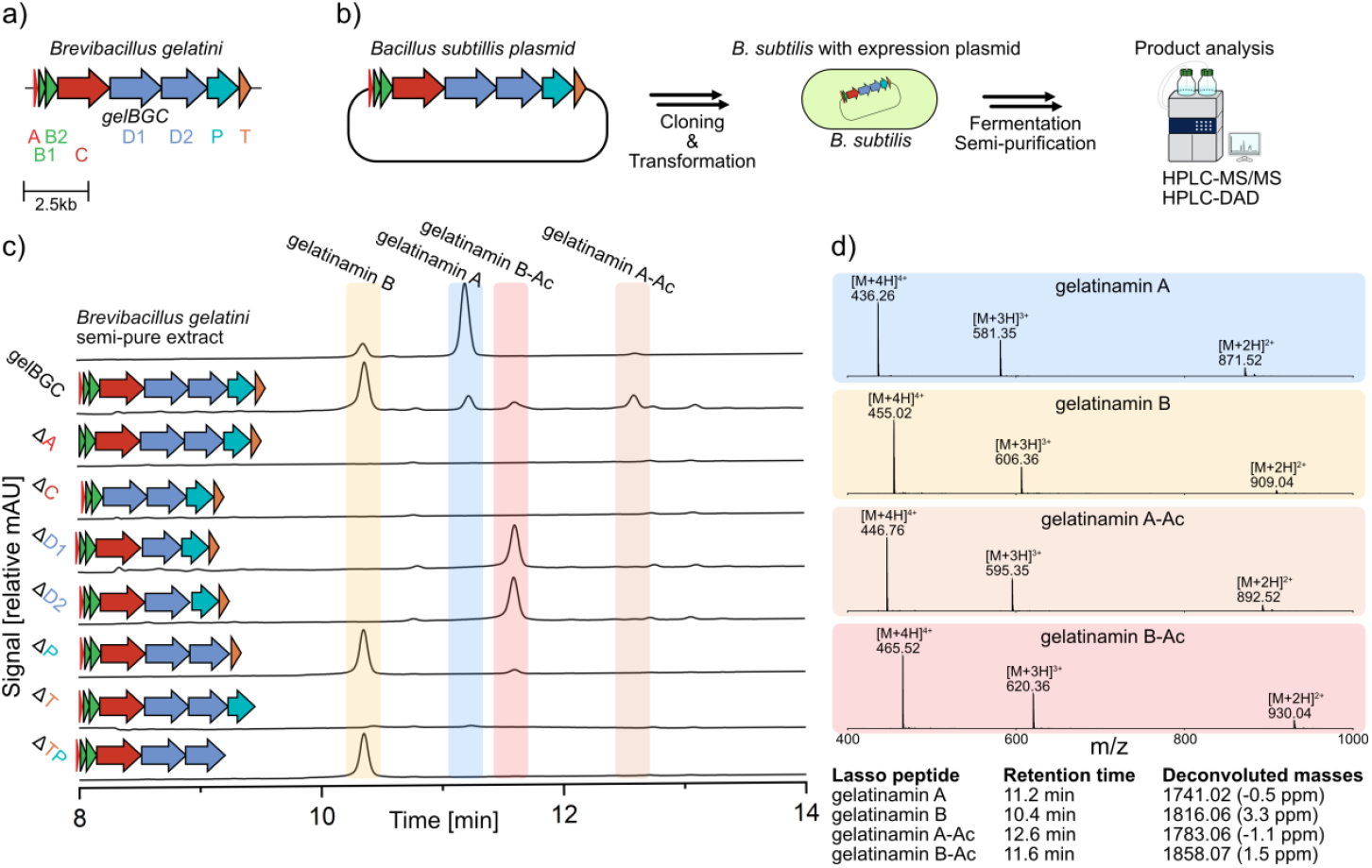
Gelatinamin biosynthesis a) The *gel* BGC found in the genome of *B. gelatini* PDF4. b) Schematic of the heterologous expression strategy applied using *B. subtilis* and analysis. c) HPLC-DAD (210 nm) chromatogram showing the semi-purified extracts from *B. gelatini* (top chromatogram) and *B. subtilis* harboring the full or partial *gel* BGC. d) MS spectra of the four dominant gelatinamin variants along with retention times and deconvoluted masses. Reference chromatograms of each purified variant and their annotated MS/MS can be found in Figure S3 and S12-S20. CAGECAT was used to visualize the BGCs.^[34]^

The extract from the supernatant of *B. gelatini* (native expression) was compared to the extract from the supernatant of *B. subtilis* SCK6 harboring a plasmid with the full *gel* BGC to evaluate gelatinamin expression in a heterologous host (Figure 1c). HPLC-DAD, HPLC-MS, and MS/MS analysis of the *B. gelatini* extract revealed four unique gelatinamin variants produced by the native host (Figure 1c, top chromatogram). Upon expression of the gel BGC in *B. subtilis* SCK6, the four major variants were observed (Figure 1c, second chromatogram). These are the expected lasso peptide (gelatinamin B), an acetylated inactive version (gelatinamin B-Ac), the double cyclized class V lasso peptide (gelatinamin A), and finally the acetylated inactive version (gelatinamin A-Ac) (Figure 1c and Figure S3). Interestingly, the predominant product produced in *B. subtilis* was gelatinamin B, suggesting that the predicted clostripain-like transpeptidase (GelP) or the pumps (GelD1 and GelD2) may be less effective in the heterologous host, compared to the natural producer, *B. gelatini*, that generates gelatinamin A as the major product.^[19]^ This result demonstrate that the *gel* BGC is sufficient for both production and export in *B. subtilis* but with minor differences in variant ratios.

To determine the role of each gene in the biosynthetic pathway, we generated deletion variants of the *gel* BGC, each lacking one gene from the cluster, and transformed them into *B. subtilis* SCK6 (Figure 1). However, we were unable to create the plasmids lacking *gelB1* and *gelB2* despite several attempts and strategies. In all cases, sequencing repeatedly revealed plasmid rearrangements and unexpected deletions, suggesting instability or recombination during cloning. All other genes were successfully deleted, allowing us to interrogate their role in gelatinamin biosynthesis.

The extracts from *B. subtilis* SCK6 harboring the generated plasmids were analyzed by HPLCDAD, HPLC-MS, and MS/MS. The chromatograms in Figure 1c demonstrate that genes *gelA* and *gelC* are both necessary for gelatinamin B production, as expected. Deleting *gelT* led to the loss of gelatinamin production in *B. subtilis*, consistent with our previous findings for triculamin, which showed the *N-*acetyltransferase, TriT, was essential for heterologous production of triculamin in *S. albus*.^[19]^ Loss of production upon deletion suggests that acetylation may prevent intracellular toxicity and indicates GelT’s role as a self-resistance mechanism. Interestingly, deleting both *gelT* and *gelP* still lead to production and export of gelatinamin B. Deleting *gelP* lead to a complete abolishment of gelatinamin A production, supporting the hypothesis that this enzyme is responsible for the cyclization between the C-terminal and the lysine sidechain. This corroborates the data presented by Wright and co-workers on lariocidin biosynthesis.^[26]^ Finally, deletion of either gene of the predicted transporters (*gelD1* and *gelD2*) leads to a shift in distribution, yielding almost exclusively the acetylated and inactive gelatinamin B-Ac. The transporters are predicted to co-fold and the requirement for both in our heterologous host supports this (Figure S10). The shift from gelatinamin B to gelatinamin B-Ac mirrors our observation for the triculamin biosynthesis and likely mitigates toxicity from intracellular accumulation.^[19]^ The combined observation aligns with the self-resistance hypothesis of GelT and also sheds light on the cellular localization of GelP. This enzyme contains a predicted lipoprotein signal peptide tag (Sec/SPII, Figure S4), which in Gram-positive bacteria anchors proteins to the extracellular leaflet of the membrane.^[32,33]^ Thus, peptides that are not exported remain intracellular, undergo acetylation, and cannot be cyclized by GelP, explaining the lack of gelatinamin A variants expressed from *gel* BGCs with a transporter knockout (Δ*gelD1* and Δ*gelD2*). These findings collectively support a self-resistance mechanism centred around GelT and support the predicted spatial localization of GelP.

### Large-scale Production and Structural Characterization of Gelatinamin A

To enable structural characterization, bioactivity assays, and *in vitro* studies, we purified gelatinamin A from the native producer *B. gelatini* PDF4 (TSB medium, 6 x 2 L, 7 days). Gelatinamin A was purified from cell-free supernatant using cation-exchange and reverse-phase chromatography yielding 42 mg.

To verify the structure previously predicted^[19]^ by MS/MS and allow for comparison with lariocidin B,^[26]^ we conducted a series of 2D and 3D NMR experiments (Table S4) to assign most ^1^H, ^13^C, and ^15^N resonances (Table S5) and determine a solution structure of gelatinamin A. The NMR sample (29.4 mg mL^-1^ in 90:10 H_2_O: D_2_O) was prepared, and all spectra were acquired at 298 K. The TOCSY spectra showed wide amide ^1^H chemical shift dispersion (6.57-9.38 ppm), indicating a well-folded structure (Figure S21).^[35]^ In addition, the NOESY experiments confirmed the two key isopeptide bonds, one between the N-terminus Ala1 and the side chain of Asp8 and one between the C-terminus Lys17 and the side chain of Lys2 (Figures S23 and S24). The Arg13, Gly14, and Val15 residues all exhibited multiple NOESY cross peaks to residues within the ring of gelatinamin A (residues 1-8), suggesting that this region of the peptide threads through the ring. Fully automated NOE cross-peak assignment and structure determination were carried out using CYANA 2.1.^[36]^ The final set of 20 models was built using 291 distance restraints (see Table S6) with a backbone RMSD of 0.62 Å ± 0.11 and a heavy-atom RMSD of 1.37 Å ± 0.22 Å. Subsequently, each model was energy-minimized using Avogadro.^[37]^ As expected, gelatinamin A adopts a right-handed lasso peptide structure with the Gly14 residue buried within the ring. A cartoon representation of the structure is presented in Figure 2a, showing the peptide sequence with the lasso and the class V-defining macrocycle in grey, the ring in turquoise, the loop in yellow, and the tail in orange. Figure 2b displays the ensemble of the 20 lowest-energy models, and Figure 2c highlights the likely positive charges (blue) and a polar cleft composed of the highly conserved PGDG motif shared by all triculamin-like peptides. As expected, the structure shares a high degree of similarity to the crystal structure of lariocidin B which was bound within the ribosome (Figure S9).^[26]^

**Figure 2.**
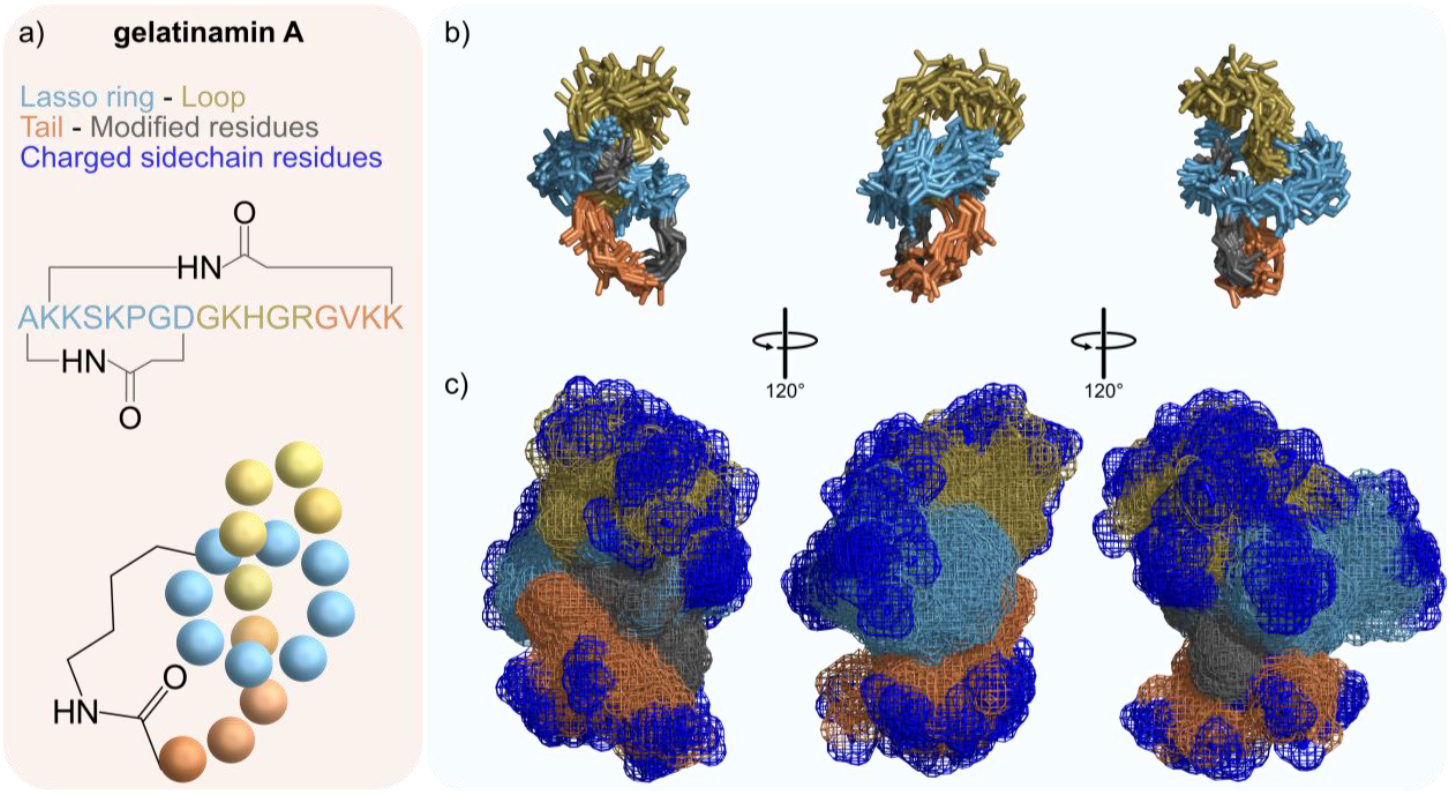
NMR-derived solution structure of gelatinamin A (PDB=9Z8J). a) Schematic representation of gelatinamin A, illustrating the two macrocycles and marking the ring (light blue), loop (gold), and tail (orange) regions of the lasso peptide. Note that one structure is shown as a non-lasso peptide for simplicity. b) Overlay of the 20 energy minimized conformers of gelatinamin A determined by NMR, showing the peptide backbone and Lys2 and Asp8 sidechains which form 2 isopeptide bonds (grey). c) All 20 individual conformers shown as sticks with sidechains and spatial mesh to provide a space-filling view of the molecule; blue indicates regions of positive charge (sidechain lysine, arginine and histidine).

### The Acetyltransferase (GelT) and Clostripain-like Peptidase (GelP) Modify Gelatinamin *in vitro*

To investigate the two tailoring enzymes encoded in the *gel* BGC, we expressed and purified GelT and GelP heterologously. Our previous work on the triculamin acetyltransferase demonstrated successful expression in *E. coli* BL21(DE3) using standard conditions,^[19]^ and although GelT shares only moderate sequence similarity with TriT (28.9 % identify, 44.6 % similarity), Alphafold3 models predict substantial structural similarity (Figure S16). The *gelT* gene was amplified from *B. gelatini* and cloned into pETM-11 using In-Fusion cloning to introduce a His_6_ tag for affinity purification. After expression and purification (Figure S5), GelT was incubated with gelatinamin A in the presence of acetyl coenzyme A (AcCoA) and the reaction was monitored using a UHPLC-DAD (Figure 3a). Emergence of a new peak (Rt = 12.5 min) only when GelT, AcCoA, and gelatinamin A were present indicated the formation of a new product with complete conversion observed after 24 h (Figure 3b). MS/MS analysis confirmed the product as gelatinamin A-Ac, matching spectra recorded on the extract from the native producer (Figure S18). Interestingly, MS/MS annotations show that GelT only acetylates Lys16 in both gelatinamin A and B (Figure S13, S18-S20), in contrast to TriT, which acetylates multiple lysine residues in gelatinamin A. ^[19]^ Importantly, in all cases examined, acetylation, irrespective of the lysine position, abolishes bioactivity.

**Figure 3.**
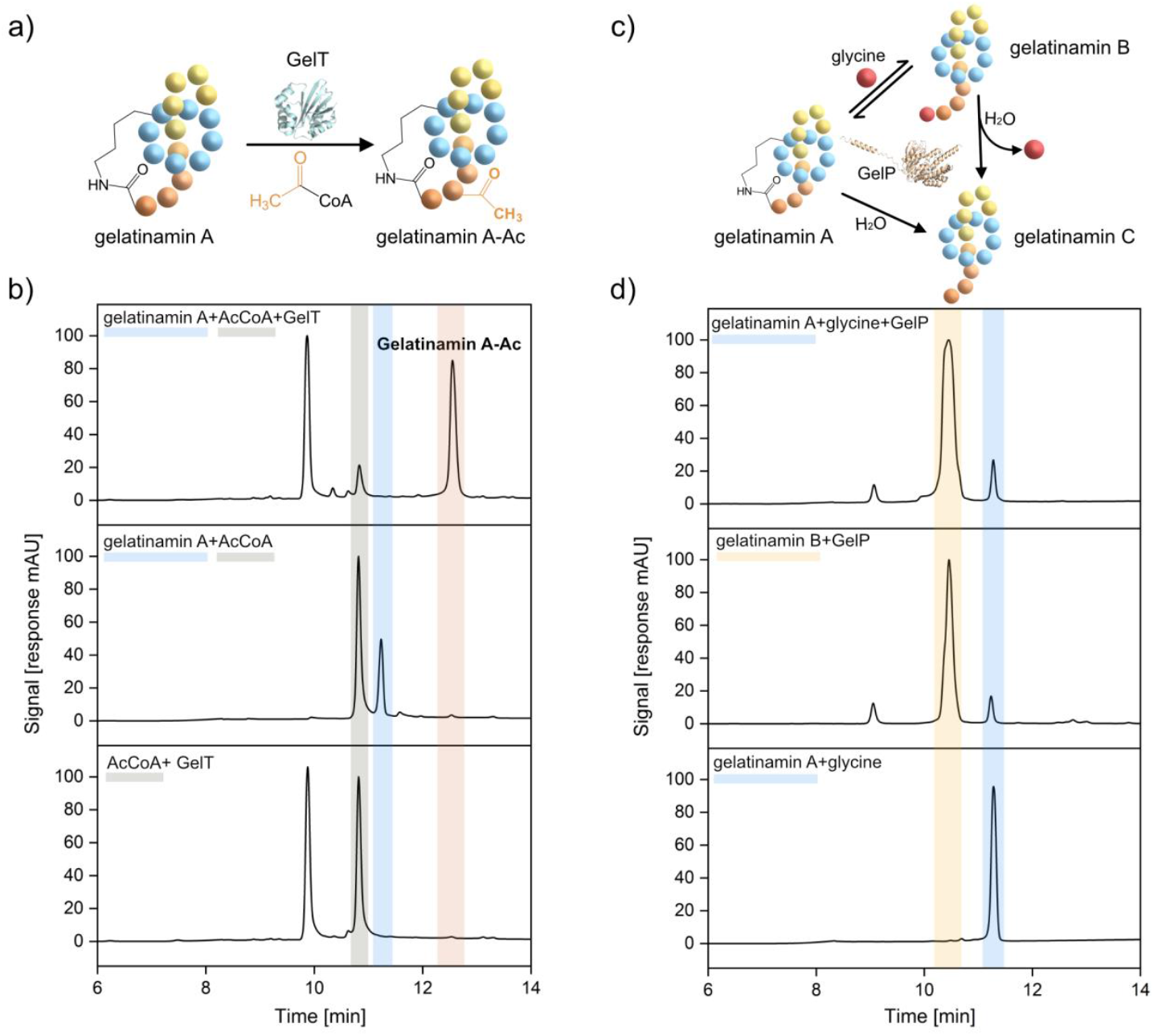
*In vitro* reaction of GelT and GelP. a) Acetylation reaction of gelatinamin A catalyzed by GelT. b) UHPLC-DAD (210 nm) chromatogram of acetylation reaction of gelatinamin A and AcCoA. The peaks are AcCoA (grey, Rt = 10.8 min), gelatinamin A (blue, Rt = 11.2 min), gelatinamin A-Ac (orange, Rt = 12.5 min). The peak at Rt = 9.9 min is hydrolyzed AcCoA (CoA) (Figure S3). c) Reversible transpeptidase reaction between gelatinamin A/B catalyzed by GelP. d) UHPLC-DAD (210 nm) chromatogram of reversible transpeptidase reactions. The top chromatogram is the GelP catalyzed transpeptidation of gelatinamin A with glycine yielding gelatinamin B (yellow, Rt = 10.4). The middle chromatogram is the GelP catalyzed intramolecular transpeptidation of gelatinamin B into A. The bottom chromatogram is gelatinamin A (blue, Rt = 11.2 min) incubated with glycine showing no turnover.

GelP is predicted to belong to the C11-like protease family and contains the conserved HisCys catalytic dyad characteristic of clostripain from *Hathewaya histolytica* and fragipain from *Bacteroides fragilis*. ^[38,39]^ Using the same strategy as for GelT, we cloned, expressed, and purified GelP with an N-terminal His_6_ tag (Figure S6). Due to an annotation error, the construct retained part of the predicted signal peptide, but since it resulted in a soluble protein in good yield, we proceeded with the *in vitro* investigation. GelP lacks the loop (Arg181-Arg188), which is removed during the calcium-dependent automaturation step known from clostripain,^[38]^ and SDS analysis showed a single-chain of ∼50 kDa, consistent with the calculated mass of 48 kDa (Figure S6). We hypothesized that the enzyme could catalyze (i) the class V defining macrocyclization converting gelatinamin B to gelatinamin A, (ii) the opening of gelatinamin A in a transpeptidase reaction using glycine, and (iii) the hydrolysis of either gelatinamin A or B, releasing gelatinamin C, which resembles gelatinamin B but lacks the Nterminal glycine due to peptide-bond hydrolysis (Figure 3c).

Reactions of gelatinamin A or B with GelP *in vitro* confirmed the transpeptidase activity of this enzyme. Incubation of gelatinamin A with GelP resulted in disappearance of gelatinamin A (blue, Rt = 11.2 min) and appearance of a peak corresponding to gelatinamin B (yellow, Rt = 10.4 min) (Figure 3d top). Conversely, incubation of gelatinamin B with GelP produced gelatinamin A (blue, Rt = 11.2 min) achieving a similar final product distribution of the two products, highlighting the microscopic reversibility of the enzyme function (Figure 3d middle). MS/MS confirmed the identities of gelatinamin A and B matching the products isolated from *B. gelatini* (Figures S15 and S17). Incubation of gelatinamin A with high concentrations of glycine (25 % w/v, ≈2800-fold molar excess to gelatinamin A) confirmed that GelP was essential to access gelatinamin B (Figure 3d bottom). Trace amounts of the hydrolysis product (gelatinamin C) were observed as a byproduct from both *in vitro* reactions by HPLC-MS/MS (Figure S14). These results confirm that GelP is a novel transpeptidase whose native function is the intramolecular transpeptidation of gelatinamin B into gelatinamin A.

### The Antibiotic Potency of Gelatinamin is Heavily Dependent on Bicarbonate

As previously described triculamin-like peptides are widely distributed across multiple phyla.^[19]^ Among these, lariocidin, has been elegantly shown by Jangra *et al*. to act as a potent ribosomal inhibitor against multiple pathogens, including clinical isolates of the ESKAPE pathogen *Acinetobacter baumannii*.^[26]^ Intriguingly, its activity is strongly dependent on culture medium. Minimal inhibitory concentrations determined in RPMI medium, which contains physiologically relevant sodium bicarbonate concentrations, revealed enhanced potency, suggesting a bicarbonate-facilitated uptake rather than transporter-mediated entry. This effect is similarly observed for macrolides and aminoglycosides when standard media are supplemented with 25 mM sodium bicarbonate.^[40–42]^.

To assess whether gelatinamin shares similar uptake characteristics, we compared its activity to lariocidin, which differs by only three positions (Ala1Ser and His11Phe, Lys17Arg). Initial MIC determinations in RPMI were unsuccessful due to poor or inconsistent bacterial growth and led us to the use of MHB-II supplemented with sodium bicarbonate. This approach allowed us to evaluate the impact of sodium bicarbonate on MIC values relative to MHB-II alone on a panel of pathogens (Table 1). The addition of bicarbonate significantly enhanced gelatinamin A activity, but the magnitude of the effect varied across species: from a modest 4-fold improvement in MIC for *E. coli* to a striking 60-fold improvement for *A. baumannii* ATCC 17978 and >200-fold improvement for a multi-drug resistant strain *A. baumannii* ATCC BAA-1605.^[28]^ Notably, the *E. coli* MC4100 *imp4213* strain, which has a permeable outer membrane, was more susceptible than the wild-type, underscoring the role of membrane permeability in peptide uptake. These results are in clear alignment with the MIC values reported for lariocidin,^[26]^ except for *Mycobacterium smegmatis*, where gelatinamin appears more potent. Together, these data suggest that bicarbonate-facilitated uptake is a general feature for the triculamin-like peptide family.

**Table 1.** MIC values for gelatinamin A tested in MHB II and MHB II supplemented with sodium bicarbonate (2 g/L ≈ 23.8 mM). All tests are performed in biological duplicates and technical triplicates. * Tested in 7H9. **Tested in 7H9 +NaHCO_3_ (2 g/L).

| Organism | MHB II | MHB II (2 g/L NaHCO<sub>3</sub>) |
| --- | --- | --- |
| *Escherichia. coli* MC4100 (WT) | 15.6 | 3.9 |
| *E. coli* MC4100 *imp4213* | 0.49 | 0.12 |
| *E. coli* MC4100 imp4213 ΔsbmA | 0.97 | $\leq 0.12$ |
| *Staphylococcus aureus* ATCC 25293 | 15.6 | 0.49 |
| *Acinetobacter baumannii* ATCC 17978 | 125 | 1.95 |
| *A. baumannii* ATCC BAA-1605 | 15.6 | 0.06 |
| *Pseudomonas aeruginosa* PA01 | >125 | 3.9 |
| *Klebsiella pneumoniae* ATCC 43816 | 125 | 15.6 |
| *Mycobacterium smegmatis* DSM 43080 | 0.03125* | 0.000488** |

Lastly, simple bioactivity assays on *M. smegmatis* revealed that acetylation abolished the antibiotic activity in similar fashion as observed for triculamin (Figure S8).^[19]^ This together with gene exclusion experiments in *B. subtilis* suggests a self-resistance mechanism.

### Hypothesized Biosynthesis of Gelatinamin A-C in *B. gelatini*

The lasso peptide gelatinamin shares considerable sequence similarity to the other reported triculamin-like peptides, triculamin,^[43]^ palmamin,^[19]^ and lariocidin^[26]^ (Figure S11), but the overall biosynthetic pathways differ in some respects. Unlike triculamin, gelatinamin, palmamin and lariocidin are from canonical lasso peptide BGCs consisting of well-characterized elements: a precursor (A), an RRE (B1), a leader peptidase (B2), a macrocyclase (C) and a predicted ATP-dependent ABC transporter(s) (Figure S10). In addition to these elements, the triculamin, gelatinamin, and lariocidin BGC contains an *N-*acetyltransferase, which conveys resistance in the case of triculamin and gelatinamin (and likely in the lariocidin case). Lastly, the gelatinamin and lariocidin BGCs harbor a clostripain-like peptidase responsible for the formation of the second cyclization between the tail and ring of the lasso peptide. Based on the explored *in vivo* biosynthesis, bioactivity and *in vitro* reconstitution of GelT and GelP, we propose the full biosynthesis of gelatinamin A/B/C (Figure 4). The pathway begins with canonical maturation of gelatinamin B: the leader sequence of GelA is recognized by GelB1, cleaved by GelB2, then folded and cyclized by GelC. Gelatinamin B is then exported to the periplasm by the GelD1D2 complex. Ineffective transport results in acetylation of gelatinamin B (Figure 1c, ΔD1, ΔD2). Once outside the cell membrane, gelatinamin B undergoes an intramolecular transpeptidase reaction catalyzed by the membrane-anchored GelP (Figure S4). Reuptake of gelatinamin A, enhanced in the presence of bicarbonate, will lead to acetylation to mitigate toxicity at high concentrations. Only trace amounts of gelatinamin C is observed as GelP has minimal peptidase activity towards gelatinamin A or gelatinamin B (Figure S14).

**Figure 4.**
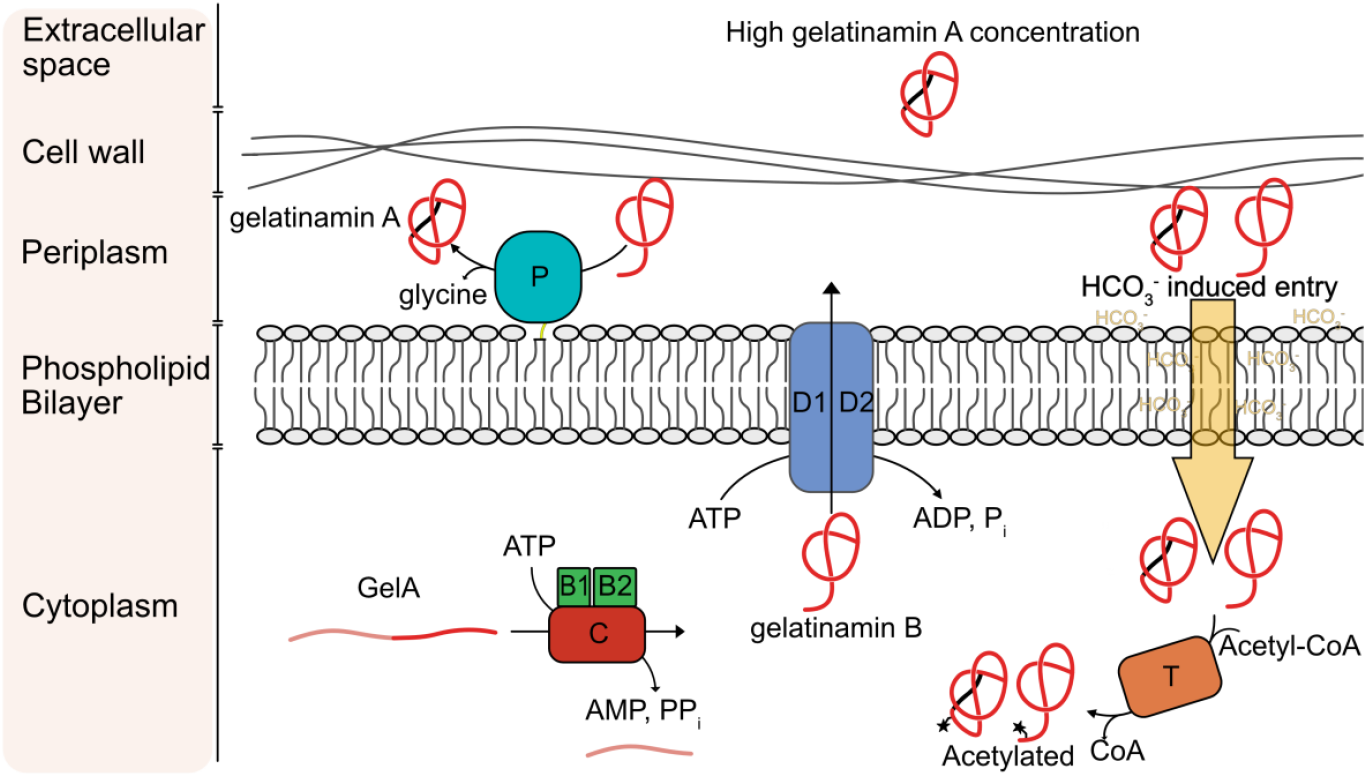
Hypothesized biosynthesis and transport of gelatinamin A-C in *B. gelatini*. Cytoplasmic biosynthesis of gelatinamin B from precursor (GelA), RRE (GelB1), leader peptidase (GelB2), macrocyclase (GelC). Export of gelatinamin B across the phospholipid bilayer by the GelD1D2 transporter complex. Maturation at the cell surface of gelatinamin B by GelP into the predominant double cyclized lasso peptide gelatinamin A. A self-resistance mechanism (GelT) acetylates bioactive gelatinamin A/B if they accumulate intracellularly.

Transport-coupled maturation is well-established for some RiPPs,^[44,45]^ but our data supports a previously unreported mechanism of periplasmic maturation by GelP (supported by Figure 1c, and Figure S4). Clostripain-like peptidases and GelP-like enzymes from lasso peptide BGCs, including LarF,^[26]^ carry predicted signal peptides, supporting this hypothesis.

## Conclusion

In this study, we expand the knowledge of triculamin-like lasso peptides, specifically gelatinamin from *B. gelatini*. We identified and analyzed the gelatinamin biosynthetic gene cluster, which closely resembles that of lariocidin and comprises genes encoding a precursor peptide (*gelA*), a RiPP recognition element (*gelB1*), a leader peptidase (*gelB2*), a lasso macrocyclase (*gelC*), two transporter genes (*gelD1* and *gelD2*), a transpeptidase (*gelP*), and an *N-*acetyltransferase (*gelT*). Using *B. subtilis* as a heterologous host, we show that gelatinamin B is produced through a canonical lasso peptide pathway (GelAB1B2C) followed by periplasmic maturation into gelatinamin A, a double cyclised figure of eight class V lasso peptide, by GelP. GelT acts as a self-resistance factor, acetylating both gelatinamin A and B at Lys16 inactivating antimicrobial activity. The absence of gelatinamin A when either transporter was deleted suggests the extracellular localization of GelP. Purification of larger quantities of gelatinamin A enabled the determination of its three-dimensional structure by NMR, and all remaining gelatinamin variants were verified by detailed MS/MS analysis. The characterization of the tailoring enzymes GelT and GelP *in vitro* confirmed that GelT-mediated acetylation parallels the resistance mechanism observed in triculamin biosynthesis. Notably, we successfully reconstituted GelP-catalyzed transpeptidation *in vitro*, demonstrating biologically relevant intramolecular cyclization of gelatinamin B to A, as well as the reversible conversion of gelatinamin A to gelatinamin B in the presence of glycine. Finally, we show that the potent bioactivity of gelatinamin A strongly depends on bicarbonate, consistent with membrane-potential–dependent uptake. Together, these results enable us to propose the complete biosynthetic pathway of gelatinamin and set the stage for further discoveries within the triculamin-like lasso peptides.

## Supporting information

Additional raw data is available at Open Science Framework (DOI:10.17605/OSF.IO/XHMRF).

## Acknowledgements

We gratefully acknowledge funding from the Carlsberg Foundation (CF22-1239) in the form of a Semper Ardens Accelerate grant [TT]. T.G.J. was supported by fellowships from the Lindemann Trust and the Omenn-Darling Bioengineering Institute. T.V. acknowledges support from the Independent Research Fund Denmark through a Sapere Aude research leader award (4251-00032B). The authors would like to thank Angela D. Zhu for initial help using Cyana 2.1.

## Data availability statement

Nuclear magnetic resonance solution structure has been deposited to the Protein Data Bank with assigned code PDB=9Z8J. Additional data is available at the Open Science Framework (DOI: 10.17605/OSF.IO/XHMRF).

