## Supporting Information for "Biosynthesis, Structure, and Antibiotic Properties of Gelatinamin A, a Triculamin-like Lasso Peptide"

[b] R. Iwase, Assist. Prof. T. Viennet, Interdisciplinary Nanoscience Center (iNANO), Aarhus University, Gustav Wieds vej 14, 8000, Aarhus, Denmark. Department of Chemistry, Aarhus University, Langelandsgade 140, 8000 Aarhus, Denmark

[c] T. G. Johnson, A. G. Zhu, Prof. A. J. Link, Department of Chemical and Biological Engineering, Princeton University, Princeton, New Jersey 08544, United States

[d] Prof. A.J. Link, Department of Chemistry, Princeton University, Princeton, New Jersey 08544, United States

[e] Prof. A. J. Link, Department of Molecular Biology, Princeton University, New Jersey 08544, United States

### General

**Data Availability**

Extended data are available on the Open Science Framework (DOI:10.17605/OSF.IO/XHMRF). The repository includes all primer sequences used for cloning and annotated plasmid files in GenBank format. In addition, all plasmids used in this study were verified by whole-plasmid sequencing (Eurofins), and the corresponding data are provided in the OSF repository. The raw NMR files are also deposited (.mnova format).

### Methods

**Nuclear Magnetic Resonance Spectroscopy**

Freeze-dried powder of gelatinamin A (14.7 mg) was dissolved in a 0.5 mL $H_2O$–$D_2O$ (90:10) solution. All NMR data were acquired at 298 K on a Bruker Neo spectrometer at a 1H frequency of 950 MHz equipped with a TCI cryoprobe and z-shielded gradients. The NMR spectra were acquired using TopSpin4.5.0 (Bruker), processed with TopSpin3.7.0, and analyzed with CcpNmr v.2.5.2.[1] as listed in Table S4. Following data acquisition, the 1H NMR chemical shifts were manually assigned (Table S5), and NOE through-space cross peaks were integrated in the 120 ms mixing time NOESY spectrum using MestReNova. 1H NMR assignments, NOE peak volumes and two explicit distance constraints defined between Ala1-Asp8 (1.33 Å between N-Cγ) and Lys2-Lys17 (1.33 Å between Nζ-C) were used for distance calibration and as upper distance restraints for structural calculations using CYANA 2.1. Seven cycles of automated NOE peak assignment and structure calculations were performed starting with 100 initial structures using 10000 torsion angle dynamic steps, to generate 20 final structures. Each of these 20 structures was then energy-minimized using the MMFF94 force field in Avogadro. The spectra and structure have been submitted to the PDB database (PDB ID: 9Z8J).

**HPLC-MS and HPLC-MS/MS**

HPLC–MS and HPLC–MS/MS measurements were carried out on an Orbitrap Exploris 120 mass spectrometer (Thermo Fisher) coupled with a Thermo Scientific SII UHPLC system. Chromatographic separation was achieved on a Luna Omega Polar C18 column (5 µm, 100 Å, 250 × 4.6 mm). The mobile phases consisted of (A) water with 0.1% formic acid and (B) acetonitrile containing 0.1% formic acid. The column temperature was maintained at 30 °C, and the flow rate was set to 1 mL/min. Samples were passed through 0.22 µm filters prior to analysis, and 10 µL was injected for each run.

The gradient program was as follows: 0–3.6 min at 0% B; 3.6–14 min, linear increase from 0% to 30% B; 14–15 min, linear increase from 30% to 95% B, followed by washing and re-equilibration of the column. MS data were collected between 3.5 and 16.5 min.

The mass spectrometer was operated using electrospray ionization in positive mode. The spray voltage was 3500 V, and the gas settings were fixed (sheath gas 60 arbitrary units, auxiliary gas 12 units, sweep gas 2 units). The ion transfer tube and vaporizer temperatures were 400 °C. Fragmentation was performed using collision-induced dissociation (CID); collision gas pressure was 1 mTorr. The Orbitrap resolution was set to 120,000 with a scan range of m/z 200–1500; the RF lens was set to 70%. Data-dependent MS/MS acquisition was enabled with minimum intensity threshold of 5000 and set resolution of 30000. HCD collision energies set to 30%. All chromatographic and mass spectrometric data were inspected and processed using Thermo FreeStyle (chromatograms, MS, deconvolution, and MS/MS spectra).

**UHPLC-DAD Analysis**

The gelatinamin products were analyzed by UHPLC (Agilent Technologies, 1260 Infinity equipped with a UV-vis detector) with a Luna Omega Polar C18 column (4.6×250 mm, 5 µm). The mobile phase used was 100% water (A) and 100% acetonitrile (B) with 0.1% trifluoroacetic acid (TFA) at a flow rate of 1 mL/min. The temperature of column was kept at 30°C. The injection volume of samples was 10 µL. The method was conducted in the following steps: 100% A was applied from 0-3.6 minutes, a linear gradient of 0-30% B was used from 3.6-14 min, a linear gradient of 30-95% B was used from 14-15.5 min, 95% B was used to wash the column for 3.5 min. In the end, the column was re-equilibrated by 100% A for 7 min. The gelatinamin compounds were detected at 210 nm.

**Production of Gelatinamin in *Brevibacillus Gelatini* PDF4**

Production and purification of gelatinamin was performed as described in our previous paper.[2] Briefly, *Brevibacillus gelatini* PDF4 (DSMZ 100115) was obtained from the German Collection of Microorganisms and Cell Cultures (DSMZ) and cultivated on TSA agar. A single colony was used to inoculate 200 mL of TSB, which was grown (37 °C, 180 rpm, 48 h). This culture was used to inoculate 6 × 2 L TSB (1:100 ratio) in baffled flasks and incubated (37 °C, 140 rpm, 5 days). After fermentation, cells were removed by centrifugation (4 °C, 12,000 rpm, 20 min). The supernatant was buffered to 50 mM HEPES (pH 8), filtered several times through a coffee filter, and stored at 4 °C until further use.

Purification was performed by cation exchange chromatography on an ÄKTA™ Start system using a 5 mL HiTrap SP XL column (Cytiva). Six purification runs were carried out, each with 2 L of culture supernatant, at a flow rate of 5 mL/min. The program used was the “short program” that is described in a later section. Fractions were analyzed for antimicrobial activity against *M. smegmatis*. Bioactive and inactive fractions were pooled separately and dialyzed against deionized water (500 Da MWCO) to remove salts.

**Separation and Purification of Gelatinamin Variants by Preparative HPLC**

Bioactive and non-bioactive pooled fractions were initially analyzed by preparative HPLC using a Luna Omega 5 µm Polar C18 100Å Column (250 x 10 mm). Initial results indicated that all variants were present in both fractions; therefore, the samples were combined for purification, lyophilized, and resuspended in 1 ml $H_2O$ 0.1% TFA. The HPLC method was subsequently optimized to achieve baseline separation of the individual variants. A flow rate of 4 mL/min and mobile phase A (0.1% TFA, $H_2O$) and mobile phase B (0.1% TFA acetonitrile) was used. 50-200 µL sample was injected per run and the method used was as follows: A multiple gradient method was used starting with 0-5% B in 10 min, from 5-25% B in 15 min, from 25-100% B in 1 min, an isocratic washing step at 100% B for 9 min, followed by an equilibration before next run. Fractions were collected in 30s time slices. Each collected fraction corresponding to a peak was freeze-dried and analyzed by HPLC-DAD and HPLC-MS. As expected, gelatinamin A was the predominant compound, followed by other derivatives in significantly lower abundance. The purified fractions were lyophilized and weighed, yielding 42 mg of pure gelatinamin A from 12L production. A representative HPLC chromatogram can be found in Figure S2.

**Heterologous Production of Gelatinamin in *Bacillus subtilis* SCK6**

<u>Cloning native Gelatinamin biosynthetic gene cluster (BGC) into shuttle plasmid with constitutive promoter</u>

The gelatinamin BGC was amplified from *B. gelatini* using PCR with primers 1 and 2 (See in OSF), which included overhangs compatible with the plasmid pMLPGB_sp_his. The plasmid was linearized by PCR using primers 3 and 4 (See Open Science Framework for full primer list). For expression of the BGC, the C-terminal $His_6$ tag was removed during plasmid linearization. The BGC and vector were assembled using In-Fusion®cloning at a 1:3 molar ratio (vector:insert). The resulting construct (*pmlpgb_sp_his_gelBGC*) was transformed into chemically competent *E. coli* DH5α cells using standard protocols and plated on LB agar containing ampicillin (100 µg/mL). After overnight incubation, single colonies were cultured in LB with ampicillin (100 µg/mL) (37 °C, 140 rpm, 18 h). Plasmids were purified using the Miniprep Kit (Thermo Fisher, K0503) and verified by whole-plasmid sequencing.

<u>Cloning native gelatinamin BGC into shuttle plasmid xylose inducible promoter</u>

To optimize production conditions, an additional construct was made exchanging the constitutive amylose promoter in pMLPGB_sp_his for an inducible xylose promoter (*xylP*) and its repressor (*xylR*). The *xylR-xylP* sequence was synthesized and ordered by Integrated DNA technologies (IDT). The previously described construct, *pmlpgb_sp_his_gelBGC* plasmid, was linearized by PCR to remove the amylase promoter, using primers 5 and 6, designed with overhangs compatible with the *xylR-xylP* fragment. The promoter and vector were assembled using In-Fusion® cloning at a 1:3 molar ratio (vector:insert). The constructs were transformed into chemically competent *E. coli* DH5α cells following standard protocols and plated on LB agar with ampicillin (100 µg/mL). After overnight incubation (37 °C), single colonies were cultured in LB with ampicillin (100 µg/mL, 37 °C, 140 rpm, 18 h). Plasmids were purified using the Miniprep Kit (Thermo Fisher, K0503) and verified by whole-plasmid sequencing.

<u>Transformation into *Bacillus subtillis* SCK6</u>

Transformation into *Bacillus subtilis* SCK6 was performed following the method described by Zhang and Zhang.[3] Briefly, *B. subtilis* SCK6 was plated on TSA containing erythromycin (3 µg/mL) and incubated overnight at 37 °C. NB: Important to use TSB/TSA without glucose to avoid carbon catabolite repression (see Table S1 for recipe). Single colonies were used to inoculate 20 mL LB supplemented with erythromycin (3 µg/mL) and incubated (37 °C, 140 rpm, 12 h). Cultures were then diluted in fresh LB containing 1% (w/v) xylose to an $OD_{600}$ of 1.0 and incubated for an additional 2 hours. The resulting cultures were either used immediately for transformation or aliquoted and stored at –80 °C with 10% (v/v) glycerol. For transformation, 500–1000 ng of purified plasmid was added to 200 µL of competent bacterial culture and incubated (37 °C, 140 rpm, 2 h). Following incubation, the entire volume was plated on LB agar supplemented with neomycin (20 ng/µL) and incubated (37 °C, 18 h).

Heterologous production of inducible promoter constructs

Before the heterologous production was initiated in *B. subtilis* SCK6 using the xylose-inducible promoter, the growth was evaluated in two different medias (LB and TB) to determine an appropriate induction optical density. *B. subtillis* SCK6 (without the plasmid) were grown in triplicates in both LB and TB media. Production cultures were started using a 1:100 inoculation from overnight starter cultures. Each variant was grown in baffled shaker flasks and incubated (37 °C, 140 rpm). Optical density (OD600) was measured every hour for each flask. The collected measurements were used to generate the growth curves shown in Figure S1. Based on the growth curves, TB-medium was selected for production due to its significantly higher final OD. Induction points were chosen at OD600 of 3-4 for expression in TB-media.

Single colonies of *B. subtilis* SCK6 harboring the plasmids of interest (*gel*BGC, ΔA, ΔC, ΔD1, ΔD2, ΔT, ΔP, ΔTP) were picked and grown in TB media supplemented with 20 µg/ml neomycin (37 °C, 180 rpm, 18 h). Each overnight culture was used to inoculate (1:100) two 50 mL TB supplemented with 20 µg/ml neomycin (37 °C, 180 rpm) in a baffled 250 mL Erlenmeyer flasks and grown to OD600 of 3-4. Heterologous expression was induced by D-xylose (1% w/v final concentration), and the cultures were grown for 3 days. The fermentation broths were clarified (4 °C, 24000 g, 60 min), and the supernatant was buffered (50 mM HEPES, pH=8). Before cationic exchange purification the supernatants were filtered through a sintered glass filter (13 µm). Each buffered supernatant was semi-purified by the short cationic exchange program described below.

Cation exchange purification

For peptide purification, two different methods were applied. Both used mobile phase A (cation exchange buffer A, Table S2) and mobile phase B (cation exchange buffer B) at a constant flow rate of 5 mL/min. Before loading the sample, the cation exchange column (HiTrap SP XL, 5 mL, Cytiva) was equilibrated with 5 column volumes (CV) of buffer A, followed by sample application.

In the short method, the column was washed with 5% B for 5 CV after sample loading, and peptides were eluted with 100% B for 5 CV. Fractions were collected in 2 mL aliquots. In the long method, the column was washed with 5 CV of buffer A and then subjected to a two-step elution gradient: 0–10% B over 10 CV, followed by 10–100% B over 20 CV. Fractions of 2 mL were collected throughout.

**Heterologous Production of GelT and GelP in *E. coli* BL21(DE3).**

For heterologous expressions of GelT and GelP, the pETM11 plasmid was used, which features an N-terminal $His_6$ tag for affinity purification. For both enzymes, the pETM11 plasmid was linearized using the same primers 7 and 8. The genes of *gelT* and *gelP* were amplified from *B. gelatini* using colony PCR. Primers (9 and 10 for *gelT* and 11 and 12 for *gelP*) included overhangs compatible with the pETM11 plasmid. The inserts and linearized vector were assembled using In-Fusion® cloning at a 1:3 molar ratio (vector:insert). Constructs were transformed into chemically competent *E. coli* DH5α cells following standard protocols and plated on LB agar with kanamycin (50 µg/mL). After overnight incubation (37 °C), single colonies were cultured in LB with kanamycin (50 µg/mL) (37 °C, 140 rpm, 18h). Plasmids were purified using the Miniprep Kit (Thermo Fisher, K0503) and verified by whole-plasmid sequencing.

Chemically competent *E. coli* BL21(DE3) cells were transformed with sequence-verified plasmids for protein expression. Single colonies were used to inoculate baffled Erlenmeyer flasks containing LB broth (100 mL) with kanamycin (50 µg/mL) and incubated (37 °C, 140 rpm, 18 h). Overnight cultures were used to inoculate large-scale production flasks (3 × 2 L LB with kanamycin, 50 µg/mL) at a 1:100 dilution. Cultures were incubated (37 °C, 140 rpm) and OD600 was monitored until it reached 0.4–0.8. At this point, cultures were cooled (ice bath, 15 min), induced with IPTG (final conc. 0.5 mM), and incubated (18 °C, 140 rpm, 18 h). Cells were harvested by centrifugation (4 °C, 8000 rpm, 20 min), and the resulting pellets were stored (-20 °C) until further use.

**Ni-NTA Affinity Purification of GelT and GelP**

Frozen cell pellets from GelT and GelP expression were thawed, weighed, and resuspended in lysis buffer (Table S2) at a ratio of 4 mL buffer per gram of pellet. The suspensions were mixed at room temperature using a magnetic stirrer for 30 minutes to initiate enzymatic lysis. Afterwards, cells were subjected to sonication on ice with an MS73 probe (60% amplitude, 5 min total, 1 s on/1 s off), and the program was repeated once to ensure thorough disruption. The resulting lysates were clarified by centrifugation (4 °C, 12,000 rpm, 20 min), and the supernatants were passed through a 0.4 µm filter prior to purification.

Protein purification was carried out on an ÄKTA™ start system equipped with a 5 mL HisTrap FF Crude column (Cytiva, 11000458) at a flow rate of 5 mL/min. The full chromatographic method, including wash and elution steps, is described in Table S3. During purification, UV absorbance was monitored continuously, and fractions showing elevated absorbance were collected and analyzed by SDS-PAGE to confirm the presence of the target protein (Figures S5 and S6).

Fractions confirmed to contain the desired protein were pooled and dialyzed against the appropriate buffer (Table S2) to remove imidazole. Final concentration of the dialyzed protein was achieved using Vivaspin 6 centrifugal filters (10 kDa MWCO, Sigma-Aldrich) to reach 1 mg/mL. Protein samples were stored at -80 °C until further use.

***In vitro* Acetylation of Gelatinamin A with GelT**

The gelatinamin A acetylation reaction was reconstituted in 50 mM Tris-HCl buffer, 10 mM $CaCl_2$, mixed with 1.5 mg/mL GelT, 2 mg/mL gelatinamin A and 5 mM acetyl-CoA (AcCoA). The reaction was incubated (37 °C, 180 rpm, 24 h). After incubation, the reaction was terminated by incubating it at 95 °C for 10 min. Control reactions were set up in parallel under identical conditions and included: (1) gelatinamin A, (2) gelatinamin A with AcCoA, and (3) GelT with AcCoA.

The products were analyzed by UHPLC (Agilent Technologies, 1260 Infinity equipped with a UV-vis detector), and HPLC-MS as described in the above sections.

**Plate Based Bioactivity Screening of *in vitro* Reactions**

Antibacterial activity was assessed against *Mycobacterium smegmatis* and *Mycobacterium phlei* using Mueller-Hinton Agar (MHA) plates. Frozen bacterial stocks (stored at -80 °C) were thawed and streaked onto fresh MHA plates, followed by overnight incubation (37 °C).

For each test, a sterile cotton swab was used to transfer culture from either *M. smegmatis* or *M. phlei* onto fresh MHA plates, creating an even lawn across the surface. Test substances (20 µL) were then spotted directly onto the inoculated plates. Plates were incubated overnight (37 °C), and initial inhibition zones were recorded the next morning. To allow clear zone definition, plates were left at room temperature for an additional 2–3 days before final evaluation.

***In vitro* Transpeptidase Reaction from Gelatinamin A to Gelatinamin B Catalyzed by GelP**

To investigate the reversible transpeptide reaction between gelatinamin A and gelatinamin B catalyzed by GelP. Prior to the assay, purified GelP was thawed from −80 °C, and was incubated with dithiothreitol (DTT, 50 mM) on ice for 2 hours. Gelatinamin A transpeptidase reaction was reconstituted in 50 mM Tris-HCl buffer (pH 7.5), 10 mM $CaCl_2$, mixed with 1 mg/mL GelP, 2 mg/mL gelatinamin A and glycine to a final glycine concentration of 25% (w/v). The reaction was incubated (37 °C, 180 rpm, 2 h). The rection was terminated by incubating it at 95 °C for 10 min. The mixture was collected by centrifugation (4 °C, 12,000 rpm, 4 min) and the precipitate was removed. Control reactions were performed in parallel under identical conditions: (1) glycine alone, (2) gelatinamin A alone, and (3) gelatinamin A with glycine (without GelP) The products were analyzed by UHPLC and HPLC-MSMS as described above.

***In vitro* Transpeptidase Reaction from Gelatinamin B to Gelatinamin A Catalyzed by GelP**

For the gelatinamin B transpeptidase reaction, the gelatinamin B compound was collected from the gelatinamin A reaction by prep-HPLC and freeze-dried to remove the solvents (as described above). The gelatinamin B transpeptidase reaction was determined in 50 mM Tris-HCl buffer (pH 7.5), 10 mM $CaCl_2$, mixed with 1 mg/mL GelP and purified gelatinamin B at 37 °C for 24 h. The reaction was terminated by incubating it at 95 °C for 10 min. The mixture was collected by centrifugation (12,000 rpm, 2 min) to remove precipitate. The products were analyzed by the same method mentioned above.

**Microbroth Dilution Assays (MIC)**

Initial testing was attempted in RPMI 1640 medium; however, growth of the strains in 96-well plates was poor, and they showed limited growth even in culture tubes. Therefore, Mueller-Hinton Broth II (MHB II) was used instead, with and without supplementation of sodium bicarbonate (2 g/L, pH-adjusted with HCl).

MICs was determined by broth microdilution (<125 µg/ml) in MHB II with or without 2 g/L $NaHCO_3$ against *Escherichia coli* MC4100, *Escherichia coli* MC4100 *imp4213*, *Escherichia coli* MC4100 imp4213 Δ*sbmA*, *Staphylococcus aureus* ATCC 25293, *Acinetobacter baumannii* ATCC 17978, *Acinetobacter baumannii* ATCC BAA-1605, *Pseudomonas aeruginosa* PA01, and *Klebsiella pneumoniae* ATCC 43816. For each strain that was tested, single colonies were used to inoculate 5 mL MHB II and incubated (37 °C, 250 rpm, 18 h). The following day, overnight cultures were diluted 1:100 into fresh MHB II and grown to mid-log phase ($OD_{600}$ = 0.4–0.5). $OD_{600}$ was measured, and cultures were diluted to $5 \times 10^6$ CFU/mL in MHB II medium, assuming an $OD_{600}$ of 1 corresponds to $1 \times 10^9$ CFU/mL. Stock solutions of gelatinamin A were prepared in 96-well plates by performing a two-fold serial dilution (11 steps). An inoculum of 20 µL ($5 \times 10^6$ CFU/mL) was added to each well using a multichannel pipette. Plates were incubated at (37 °C, 250 rpm, 18 h). The following morning, the ODs were measured with a plate reader.

For *Mycobacterium smegmatis* DSM 43080 MIC was determined by broth microdilution (<32 µg/ml) in Middlebrook 7H9 broth with 0.4% glycerol and 10% ADC (0.5% BSA, 0.2% dextrose, and 0.003 g catalase/L) with or without 2 g/L $NaHCO_3$. The inoculum was standardized to an approximate concentration of 5 × $10^5$ CFU/ml. Plates were incubated at 37 °C with shaking at 250 rpm for 18h. Mycobacterial growth was assessed after 2-5 days at 37 °C. MIC was defined as the lowest concentration of compound with no visible growth. Experiments were performed with technical triplicates and two biological replicates.

**Table S1.** List of media recipes.

| Name | Composition |
|---|---|
| Mueller Hinton broth (MHB) | Dextrose 1.0 g/L, tryptone 5.0 g/L, yeast extract 2.5 g/L |
| Mueller Hinton broth – II (MHB II) | Dextrose 1.0 g/L, tryptone 5.0 g/L, yeast extract 2.5 g/L, calcium 20-25 mg/l, magnesium 10-12.5 mg/l. |
| Mueller Hinton Agar (MHA) | Dextrose 1.0 g/L, tryptone 5.0 g/L, yeast extract 2.5 g/L, agar 9.0 g/L |
| LB Broth | Tryptone 10.0 g/L, yeast extract 5.0 g/L, NaCl 10.0 g/L |
| LB Agar | Tryptone 10.0 g/L, yeast extract 5.0 g/L, NaCl 10.0 g/L, agar 15.0 g/L |
| Tryptic soy broth (TSB) | Peptone from casein 17.0 g/L, peptone from soymeal 3.0 g/L, NaCl 5.0 g/L, di-potassium hydrogen phosphate 2.5 g/L |
| Tryptic soy agar (TSA) | Peptone from casein 17.0 g/L, peptone from soymeal 3.0 g/L, NaCl 5.0 g/L, di-potassium hydrogen phosphate 2.5 g/L, agar 15 g/L |
| Terrific Broth (TB media) | Tryptone 12.0 g/L, yeast extract 24.0 g/L, $KH_2PO_4$ 2.31 g/L, $K_2HPO_4$ 12.54 g/L, glycerol 0.4% v/v. |
| Middlebrook 7H9 broth with ADC | Ammonium sulfate 0.50 g/L, disodium phosphate 2.50 g/L, monopotassium phosphate 1.00 g/L, sodium citrate 0.10 g/L, magnesium sulfate 0.05 g/L, calcium chloride 0.0005 g/L, zinc sulfate 0.001 g/L, copper sulfate 0.001 g/L, ferric ammonium citrate 0.04 g/L, L-glutamic acid 0.50 g/L, pyridoxine 0.001 g/L, biotin 0.0005 g/L, plus 0.5% BSA, 0.2% dextrose, 0.003 g catalase/L, glycerol 4 ml/L, with or without 2 g/L $NaHCO_3$. |

**Table S2.** List of buffer concentrations.

| Name | Composition |
|---|---|
| Lysis Buffer | 20 mM phosphate buffer (pH 7.4), 200 mM NaCl, 0.2 mg/ml lysozyme, 20 µg/ml DNAse, 1 mM MgCl2 |
| Dialysis buffer | 20 mM phosphate buffer (pH 7.4), 200 mM NaCl |

| | |
|---|---|
| Cation exchange buffer A | 50 mM HEPES (pH 8) |
| Cation exchange buffer B | 50 mM HEPES (pH 8), 1M NaCl |
| His-tag purification buffer A | 20 mM phosphate (pH 7.4), 500 mM NaCl, 10 mM imidazole |
| His-tag purification buffer B | 20 mM phosphate (pH 7.4), 500 mM NaCl, 500 mM imidazole |

**Table S3.** AKTA program for his-tag purification of GelT and GelP

| **Column** | **5 mL HisTrap FF crude** | |
|---|---|---|
| **Flowrate** | **5 mL/min** | |
| **Steps** | **Action** | **Duration (CV)** |
| 1 | Equilibrating column in water | 2 |
| 2 | Equilibration of column in buffer A | 5 |
| 3 | Sample application | - |
| 4 | Column wash – wash unbound sample (buffer A) | 10 |
| 5 | Elution, gradient (0% to 25% buffer B) | 10 |
| 6 | Elution, isocratic (100% buffer B) | 10 |
| 7 | Equilibration in buffer A | 5 |

### Results

**Evaluation of Growth of Heterologous Host *Bacillus subtilis* SCK6**

To determine suitable growth conditions for heterologous expression, we evaluated the growth performance of *Bacillus subtilis* SCK6 in LB and TB media. Cultures were incubated (37 °C, 140 rpm), and optical density ($OD_{600}$) was monitored over time in biological triplicates. The purpose was to identify the medium supporting the highest final cell density and to determine the appropriate induction point. *Bacillus subtilis* SCK6 reached higher final ODs in TB compared to LB, indicating that TB provides a richer nutrient environment supporting more robust growth. Based on the obtained growth curves, mid-log phase ($OD_{600} \approx 3–4$) was selected as the optimal time for induction in subsequent expression experiments in TB.

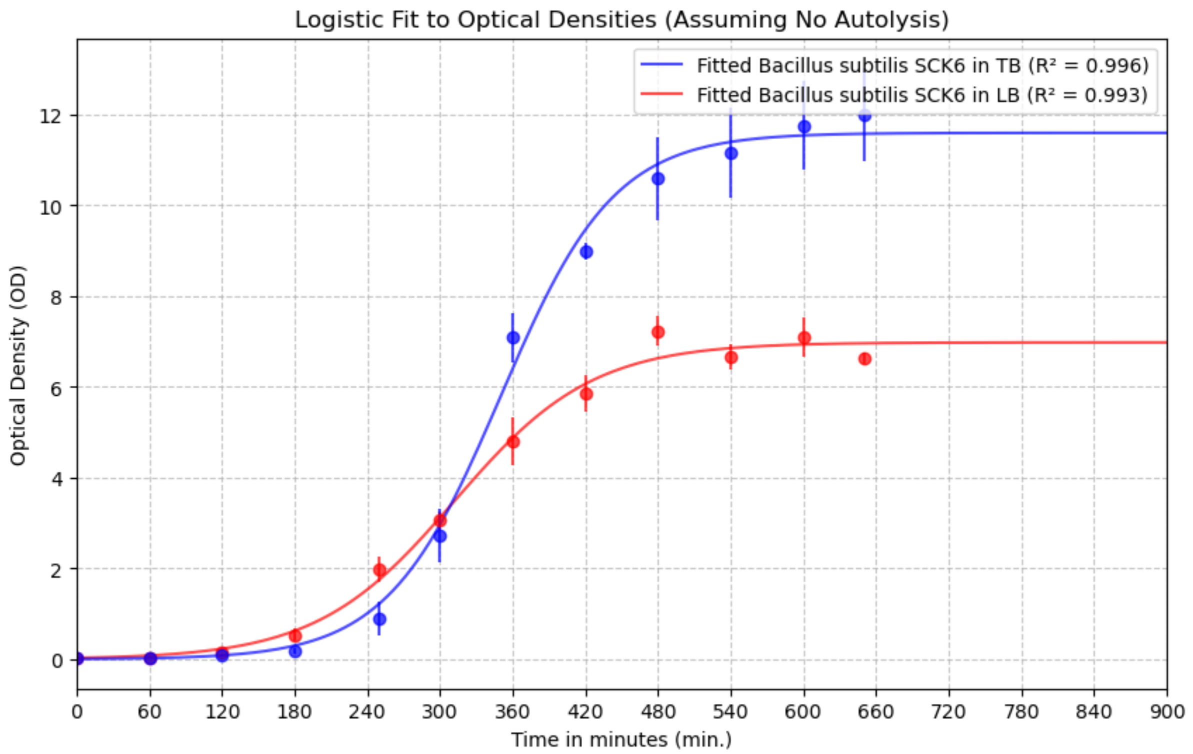


**Figure S1.** Growth curves of *Bacillus subtilis* SCK6 strains cultivated in LB and TB media. Optical density at 600 nm ($OD_{600}$) was measured over time, and a logistic growth model was fitted to the data (assuming no autolysis). The two curves represent *B. subtilis* SCK6 strains grown in LB medium (red) and in TB medium (blue). The strain reached higher final optical densities in TB compared to LB, indicating enhanced biomass accumulation in the richer medium. Error bars represent standard deviations from biological triplicates.

**Preparative HPLC-UV Purification of Gelatinamin A from *Brevibacillus gelatini***

A representative preparative HPLC-UV chromatogram from the purification of gelatinamin produced by the native producer, *Brevibacillus gelatini* is shown in Figure S2. Gelatinamin A was identified as the major product and was collected based on its retention time. The pooled fractions were lyophilized and subsequently resuspended in water, which was repeated twice to remove residual trifluoroacetic acid (TFA).

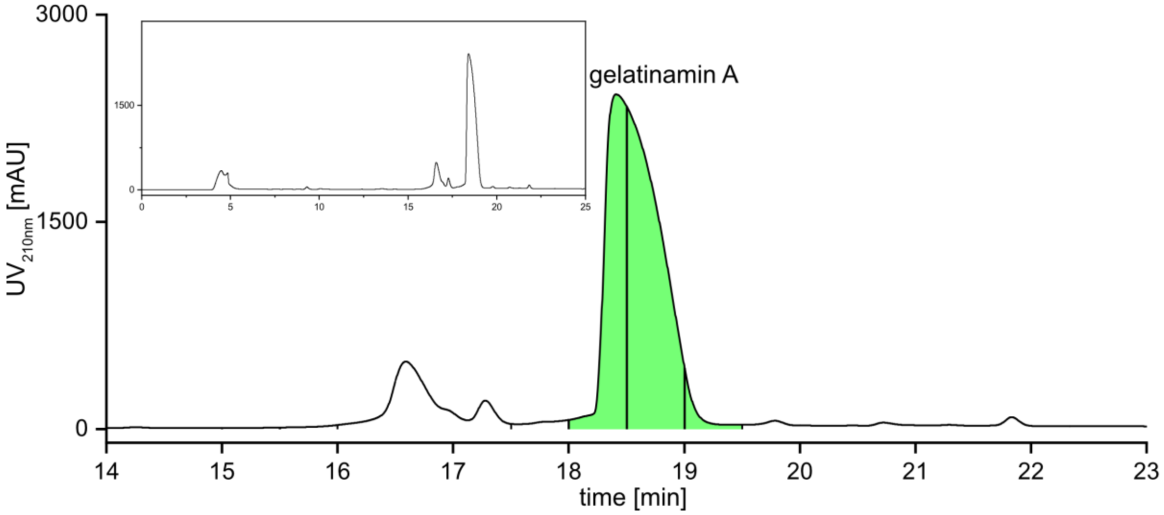


**Figure S2.** Preparative HPLC-UV chromatogram of the gelatinamin A purification from *Brevibacillus gelatini*. Vertical lines indicate 30-second time-slice fractionation. The full chromatogram is shown in the top left corner.

**UHPLC Standard Chromatograms**

Representative UHPLC chromatograms of the purified standards used for reference are shown in Figure S3. Gelatinamin A standard was purified from *Brevibacillus gelatini* PDF4 production. Gelatinamin B, gelatinamin A-Ac**,** gelatinamin B-Ac standards were purified from reactions of gelatinamin A transpeptidation, gelatinamin A acetylation and gelatinamin B acetylation, respectively. AcCoA and CoA were from commercial products (Sigma-Aldrich). All the gelatinamin standards were analyzed under identical conditions and further confirmed by HPLC-MS/MS. The compounds eluted with distinct retention times at 11.2, 10.4, 12.6, 11.6, 10.8, and 9.9 minutes, respectively, allowing unambiguous identification in subsequent assays. Relative response values were normalized to the highest signal intensity.

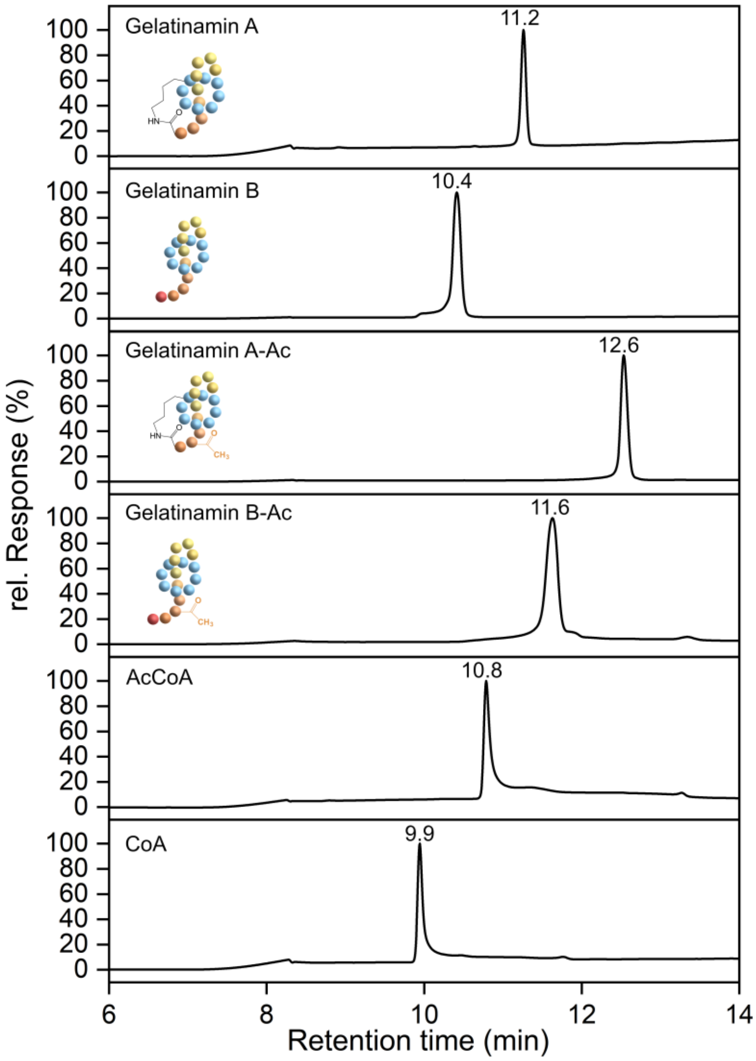


**Figure S3.** UHPLC-DAD (210nm) chromatograms of standard compounds of gelatinamin A, gelatinamin B, gelatinamin A-Ac, gelatinamin B-Ac, AcCoA and CoA.

**Signal Peptide Prediction for GelP**

To predict whether GelP is secreted via the Sec or Tat pathway, its amino acid sequence was analyzed using SignalP 6.0.[4] The prediction identified a 29-amino acid N-terminal signal peptide (MRNYTRCRLRSMCKALMMTVLAALLLVGC) characteristic of a Sec/SPII-type lipoprotein signal peptide. The analysis yielded a high likelihood (0.9998) for the Sec/SPII classification. The predicted cleavage site was located between Gly28 and Cys29, consistent with processing a type II signal peptidase. These findings suggest that GelP is a lipoprotein exported via the Sec pathway and anchored to the membrane by lipidation at the conserved cysteine residue.

a)

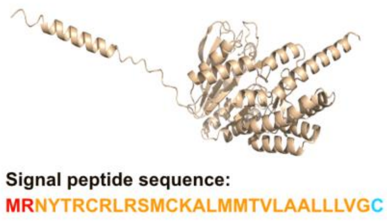


b)

| Protein type | Other | Signal Peptide (Sec/SPI) | Lipoprotein signal peptide (Sec/SPII) | TAT signal peptide (Tat/SPI) | TAT Lipoprotein signal peptide (Tat/SPII) | Pilin-like signal peptide (Sec/SPIII) |
|---|---|---|---|---|---|---|
| Likelihood | 0 | 0.0002 | 0.9998 | 0 | 0 | 0 |

c)

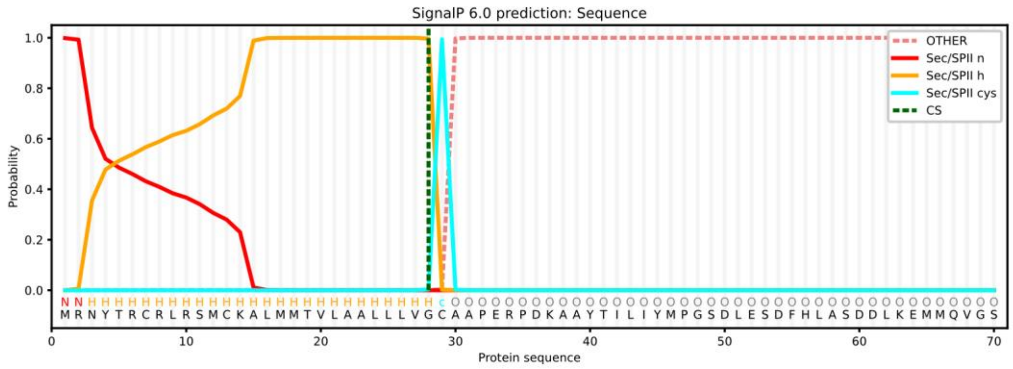


**Figure S4.** Signal peptide prediction using GelP as input. a) Predicted structure of GelP with signal peptide consisting of 29 amino acids by AlphaFold3.[5] b) Possibilities of signal peptide types for GelP predicted by SignalP 6[6]. c) Prediction of the GelP signal peptide sequence matching the Sec/SPII type of signal peptide by SignalP 6. Sec/SPII "N" refers to a positively charged amino terminal; Sec/SPII "H" refers to a hydrophobic core; Sec/SPII "C" is the required cysteine for a lipid modification; "CS" refers to the predicted cleavage site between Gly28 and Cys29 by a type II lipoprotein signal peptidase.[7]

**Purification of GelT and GelP Enzymes**

The acetyltransferase GelT and the costripain-like enzyme GelP were heterologously expressed in *E. coli* BL21(DE3) as N-terminal His-tagged proteins. Expression was induced with IPTG during mid-log phase growth in LB medium, and cells were harvested and lysed by sonication. The enzymes were purified using Ni-NTA affinity chromatography on ÄKTA purification system (Figures S5a and S6a).

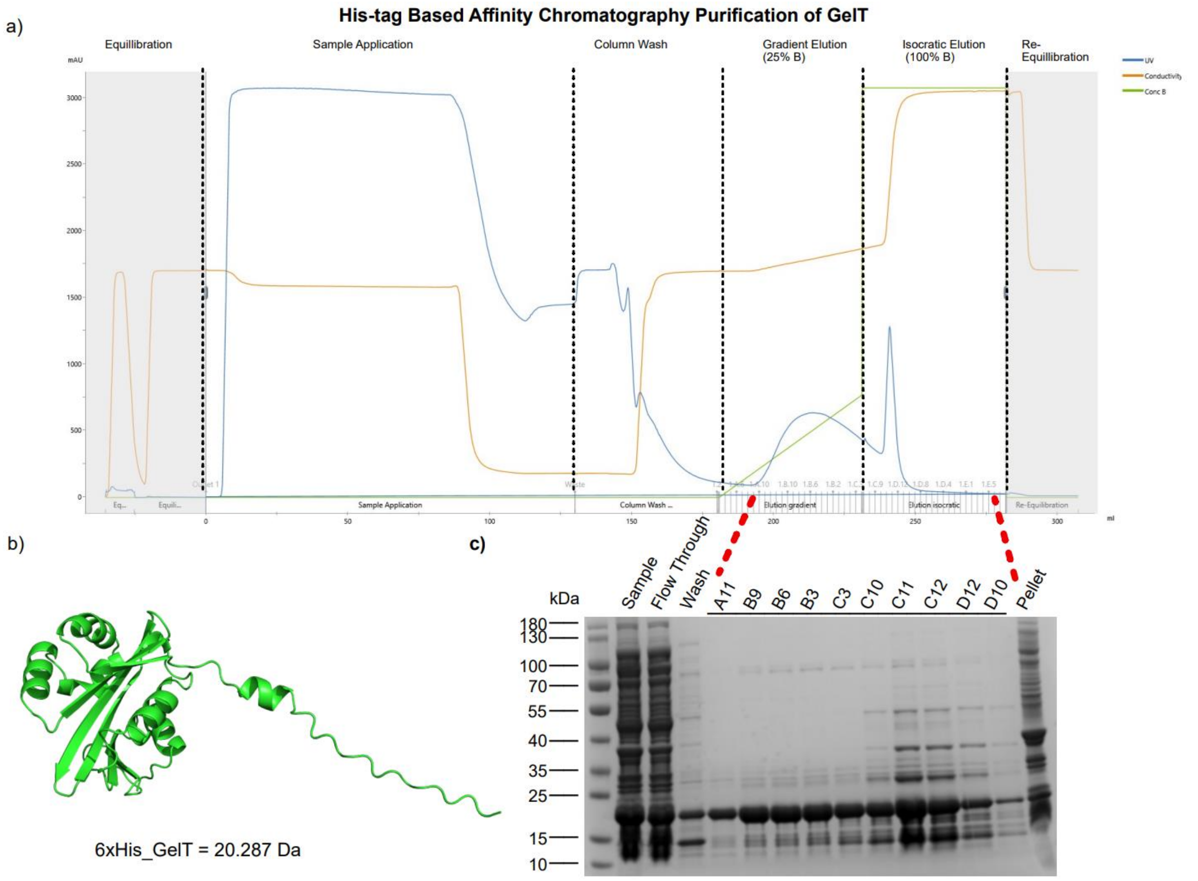


**Figure S5.** Purification and analysis of the acetyltransferase GelT. (a) Chromatogram from His-tag-based affinity purification of GelT using Ni-NTA resin. The UV absorbance (blue) and conductivity (orange) profiles are shown, with key purification stages indicated: equilibration, sample application, column wash, gradient elution (25% buffer B), isocratic elution (100% buffer B), and re-equilibration. Fractions collected during the elution phase (A11–D12) were analyzed by SDS–PAGE. (b) Predicted 3D Alphafold3[5] structure of GelT. (c) SDS–PAGE analysis of purification fractions showing GelT enrichment in the elution fractions (C10–D12). Lane labels correspond to sample processing steps and collect fractions.

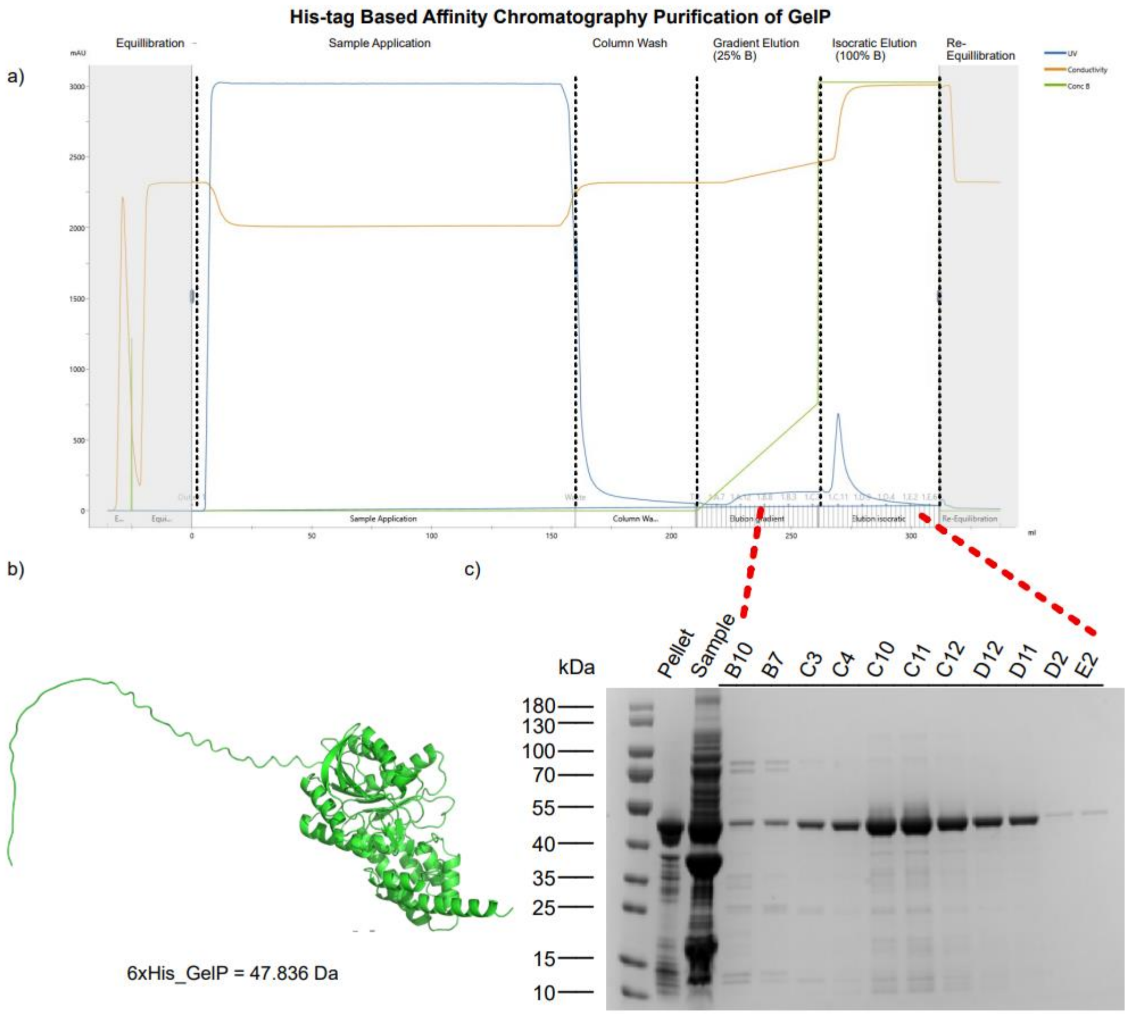


**Figure S6.** Purification and analysis of the transpeptidase GelP. (a) Chromatogram from His-tag-based affinity purification of GelP using Ni-NTA resin. The UV absorbance (blue) and conductivity (orange) traces are shown, with distinct purification steps indicated: equilibration, sample application, column wash, gradient elution (25% buffer B), isocratic elution (100% buffer B), and re-equilibration. Fractions collected during the elution phase (B7–E2) were analyzed by SDS–PAGE. (b) Predicted 3D Alphafold3[5] structure of GelP. (c) SDS–PAGE analysis of purification fractions showing GelP enrichment in the elution fractions (C4–D12). Lane labels correspond to the different purification fractions collected during the affinity chromatography process.

**Comparison of GelT to Triculamin *N*-acetyltransferase (TriT)**

Both TriT and GelT are *N*-acetyltransferases that convey resistance. TriT acetylates a lysine residue in the ring of triculamin, whereas GelT acetylates Lys16 of gelatinamin A and B. Alphafold3[5] structural predictions allow a structural comparison (Figure S7), which indicates structural similarities despite major size differences. Furthermore, the sequence of alignment reveals short alignment length, low identity, and moderate similarity.

a)

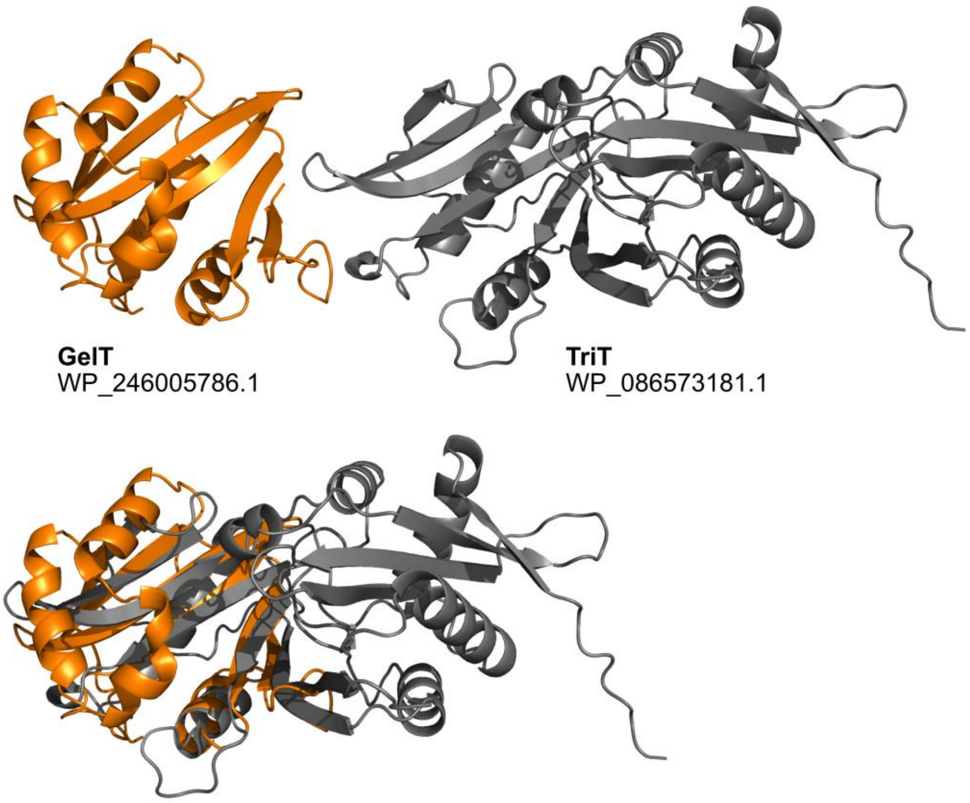


b)

**Alignment parameters:**
Smith-Waterman, Blosum62

Alignment length: 83
Identity 28.9%

Similarity: 44.6%
Gaps: 20.5%

```
GelT 70  VAMVEIEEGMEGSIGLI----VNPSLRHRGYGKTLLREVLEHEELSGVKRWIA----------GIEEDNVACLRCFEAVGFEK 138
         VA   I E ++G +G++    V P  R +G G+ LL   L  + L+G+    A          G E D  A    +E+ GF +
TriT 203 VAEAVIGEPVQG-VGVLWWIEVGPEARRQGLGRALLGSAL--DTLTGLGATEAILYVDDDEPPGGERDRTAANALYESAGFNE 282
```

**Figure S7.** a) Alphafold3[5]predicted structures of the *N*-acetyltransferase GelT and the triculamin *N*-acetyltransferase TriT. b) Alignment of GelT and TriT showing short sequence alignment, low sequence identity, and moderate similarity.

#### Spot-on-lawn of *in vitro* Acetylated Gelatinamin A using GelT

To showcase that acetylated gelatinamin A is biologically inactive, the reaction mixture with and without GelT was tested against *M. smegmatis and M. phlei*.

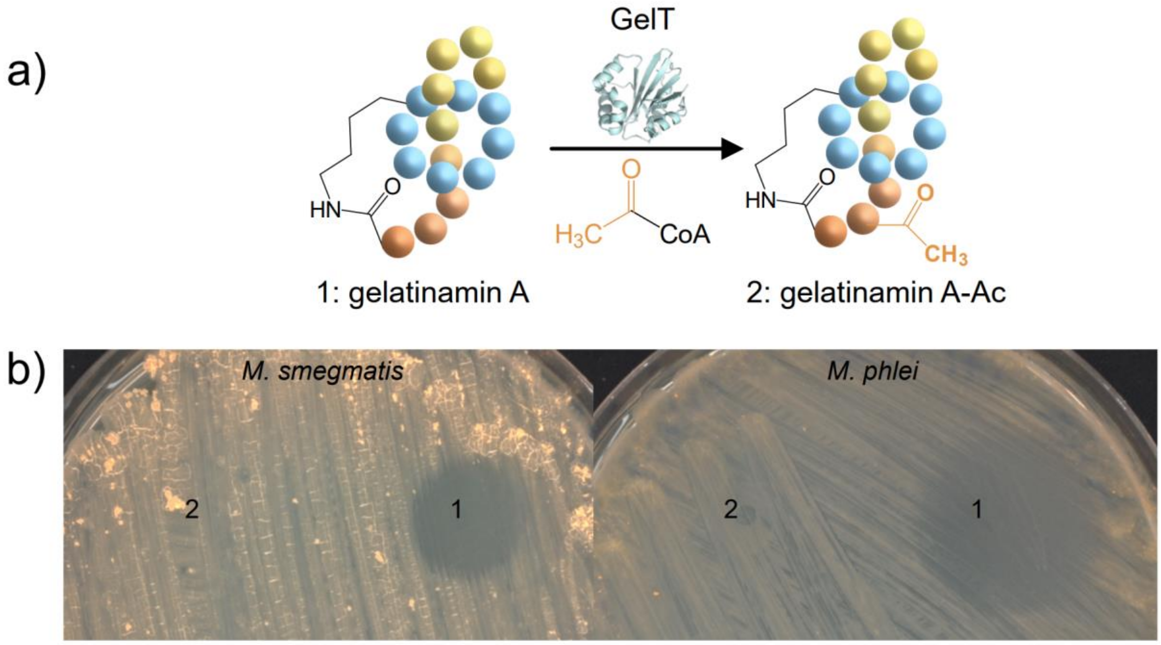


**Figure S8.** *In vitro* acetylation of gelatinamin A and spot-on-lawn bioactivity assays. a) Schematic representation of the enzymatic acetylation of gelatinamin A (1) by the acetyltransferase GelT using Ac-CoA as the donor, yielding acetylated gelatinamin A-Ac(2) b) Spot-on-lawn assays comparing the antibacterial activity of gelatinamin A (1) and gelatinamin A-Ac (2) against *Mycobacterium smegmatis* (left) and *Mycobacterium phlei* (right). A total of 10 µL of each reaction mixture was spotted onto the bacterial lawn. Numbers correspond to the peptide applied at each spot and clearly indicate complete loss of bioactivity upon acetylation.

**Comparison of Gelatinamin A to Lariocidin B (LAR-B)**

The X-ray crystal structure of lariocidin B was recently solved in complex with the ribosome of *Thermus thermophilus* (PDB=9DFD). Both gelatinamin A and lariocidin B have unprecedented Lys2 to C-terminus isopeptide bond, thus we compared the structure of gelatinamin A and lariocidin B (Figure S9).

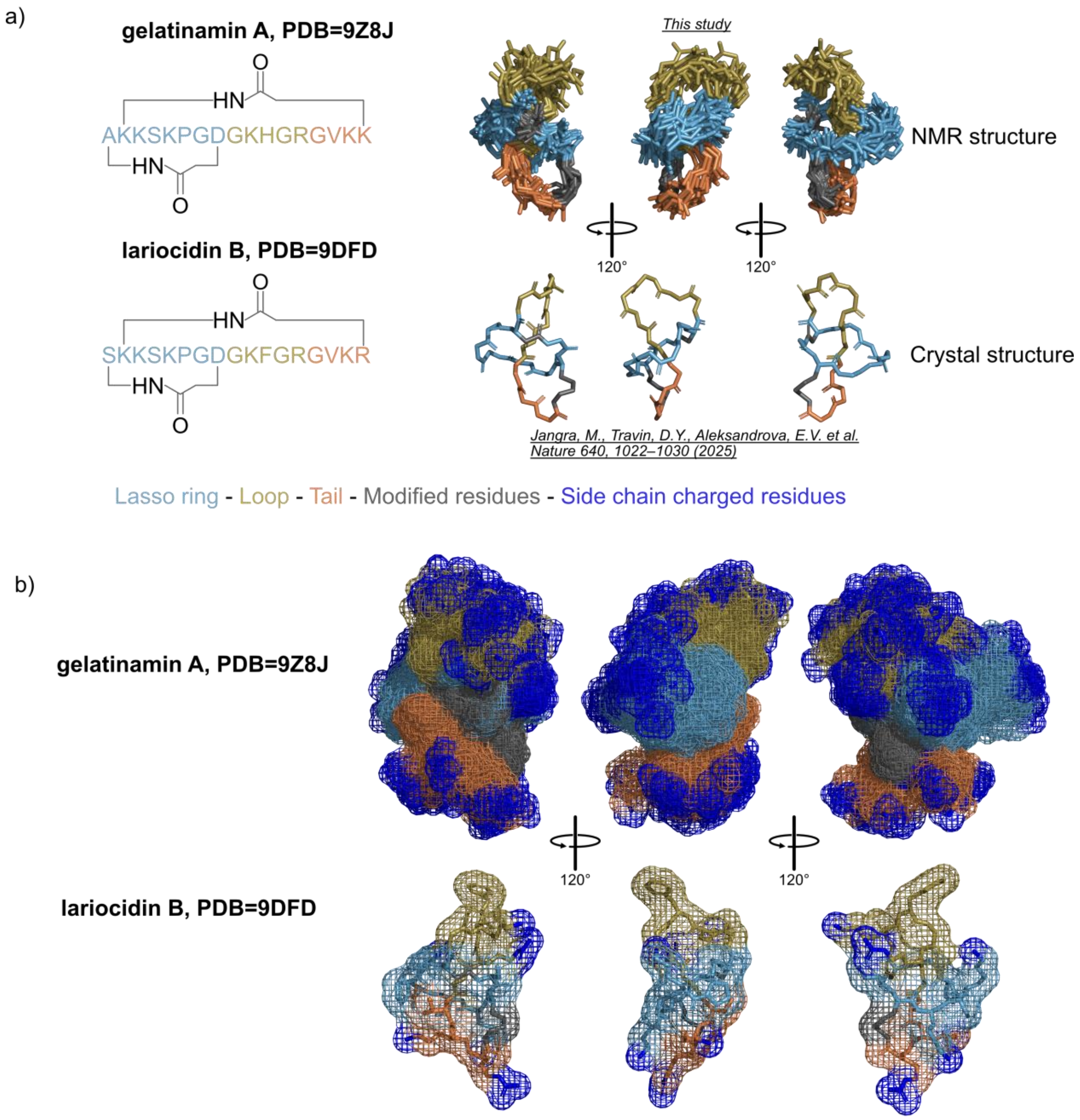


**Figure S9.** Structural comparison of gelatinamin A to lariocidin B a) Schematic representations of gelatinamin A and lariocidin B, highlighting their two macrocycles and the ring, loop, and tail regions characteristic of lasso peptides. The structural comparison includes an overlay of the 20 lowest-energy NMR conformers of gelatinamin A with the crystal structure of lariocidin B. b) All 20 individual conformers of gelatinamin A shown with their 3D spatial charge distribution, compared to the crystal structure of lariocidin B. Blue indicates regions of positive charge (sidechain lysine, arginine and histidine).

**Alphafold3 Prediction of GelD1D2 Complex with ATP and $Mg^{2+}$**

Our biosynthetic studies of GelD1 and GelD2 indicate that they both are essential for gelatinamin B export. The Alphafold3[5]a predicted structure of GelD1D2 gives reasonable ipTM score, pLDDT and PAE (Figure S10).

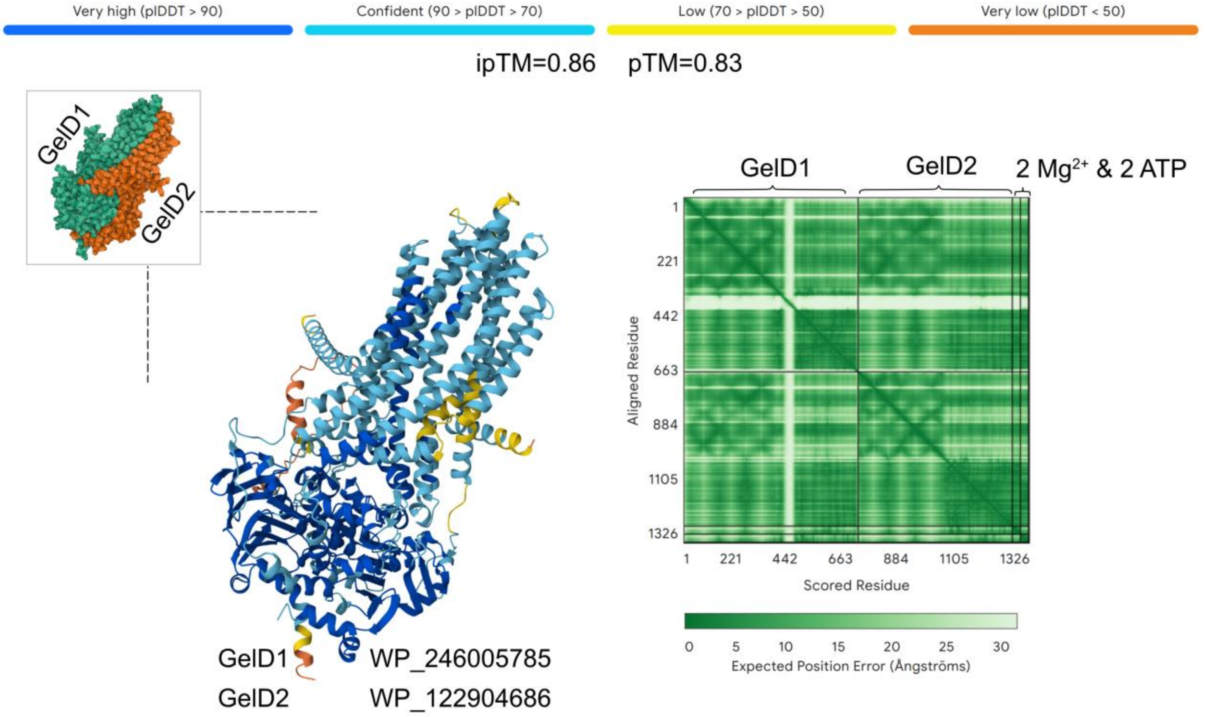


**Figure S10.** Alphafold3[5] predicted the complex of GelD1 and GelD2, which indicates that the GelD1 and GelD2 forms a complex.

**Precursor Comparison of Triculamin-like peptides, Precursor Sequences, and BGCs.**

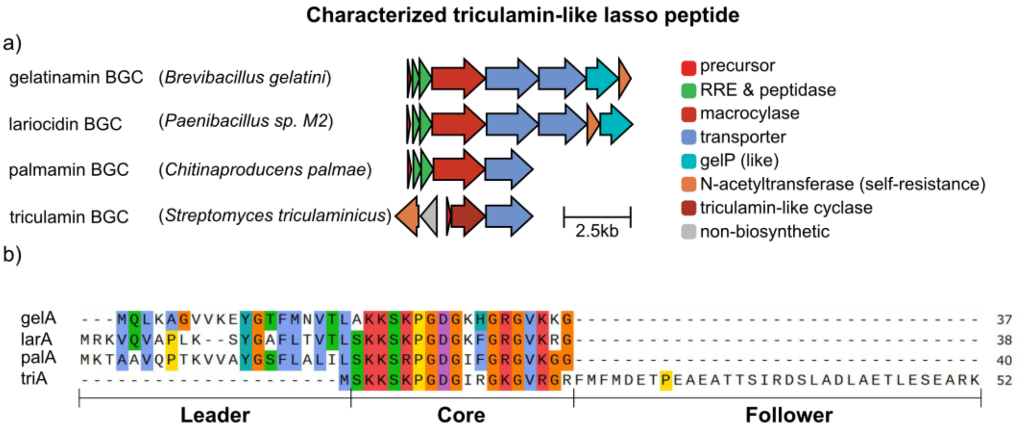


**Figure S11**. a) BGCs characterized triculamin-like lasso peptides. b) Precursors of characterized triculamin-like lasso peptides. Core sequences are highly conserved across canonical and non-canonical precursors. Note that the macrocyclases and *N*-acetyltransferases from canonical BGCs (gelatinamin, lariocidin, palmamin) and non-canonical (triculamin) are significantly different.

**MS/MS Spectra of Gelatinamin Variants: A, A-Ac, B, B-Ac, C.**

Fragmentation of lasso peptides has become an increasingly important tool for determining exotic PTMs. For lasso peptides containing big loops topological information may be gained from interlocked fragments as a result of double peptide fragments.[8] Below we present MS/MS spectra of gelatinamin A (from native producer, heterologous producer and *in vitro* GelP produced), gelatinamin A-Ac (from native producer, heterologous producer and *in vitro* GelT produced), gelatinamin B (native producer, heterologous producer and *in vitro* produced GelP), gelatinamin B-Ac (native producer, heterologous producer and *in vitro* produced) and gelatinamin C (*in vitro* produced GelP).

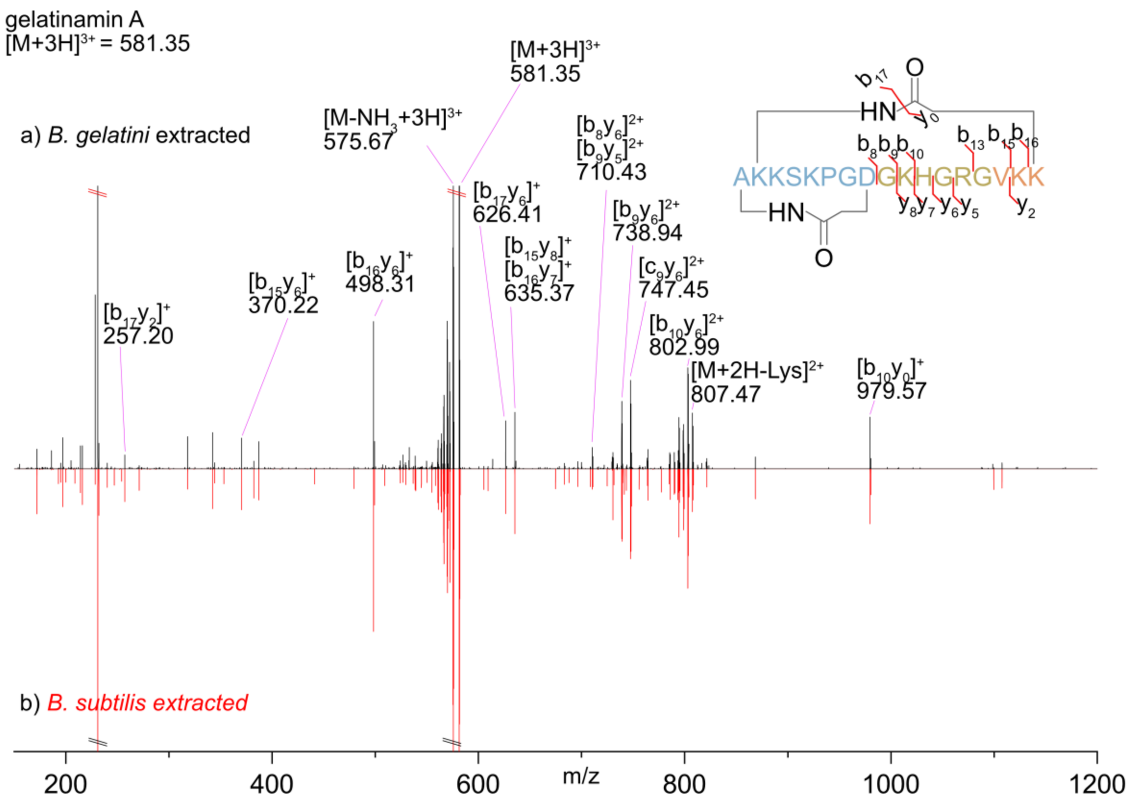


**Figure S12.** Comparison of MS/MS spectra of gelatinamin A ([M+3H]$^{3+}$]) obtained from the native producer *B. gelatini* and from recombinant expression in *B. subtilis* SCK6. The absence of both N- and C-terminus results in observation of only double fragments. a) *B. gelatini* produced gelatinamin A MS/MS spectrum. b) *B. subtilis* produced gelatinamin A MS/MS spectrum. The shown gelatinamin A sequence is not shown as a lasso peptide for simplicity.

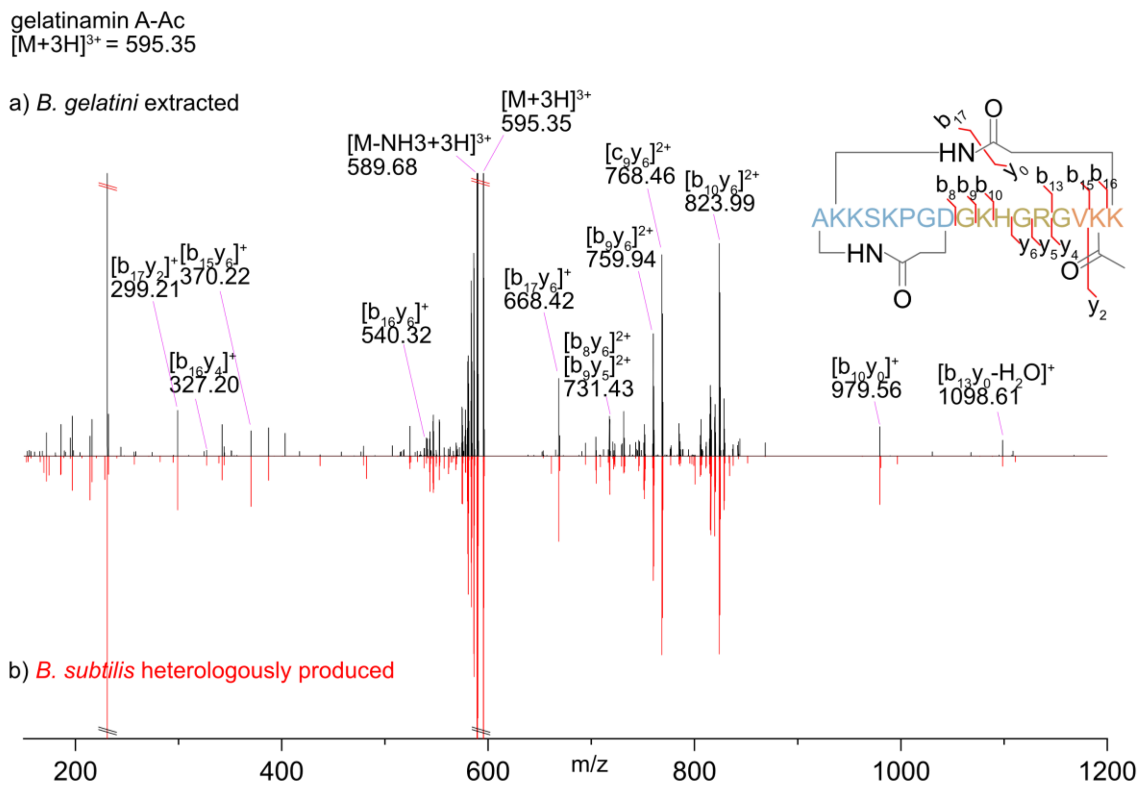


**Figure S13.** Comparison of MS/MS spectra of gelatinamin A-Ac ($[M+3H]^{3+}$) obtained from the native producer *B. gelatini* and from recombinant expression in *B. subtilis* SCK6. The absence of both N- and C-terminus results in observation of only double fragments. a) *B. gelatini* produced gelatinamin A-Ac MS/MS spectrum. b) *B. subtilis* produced gelatinamin A-Ac MS/MS spectrum. The shown gelatinamin A-Ac sequence is not shown as a lasso peptide for simplicity.

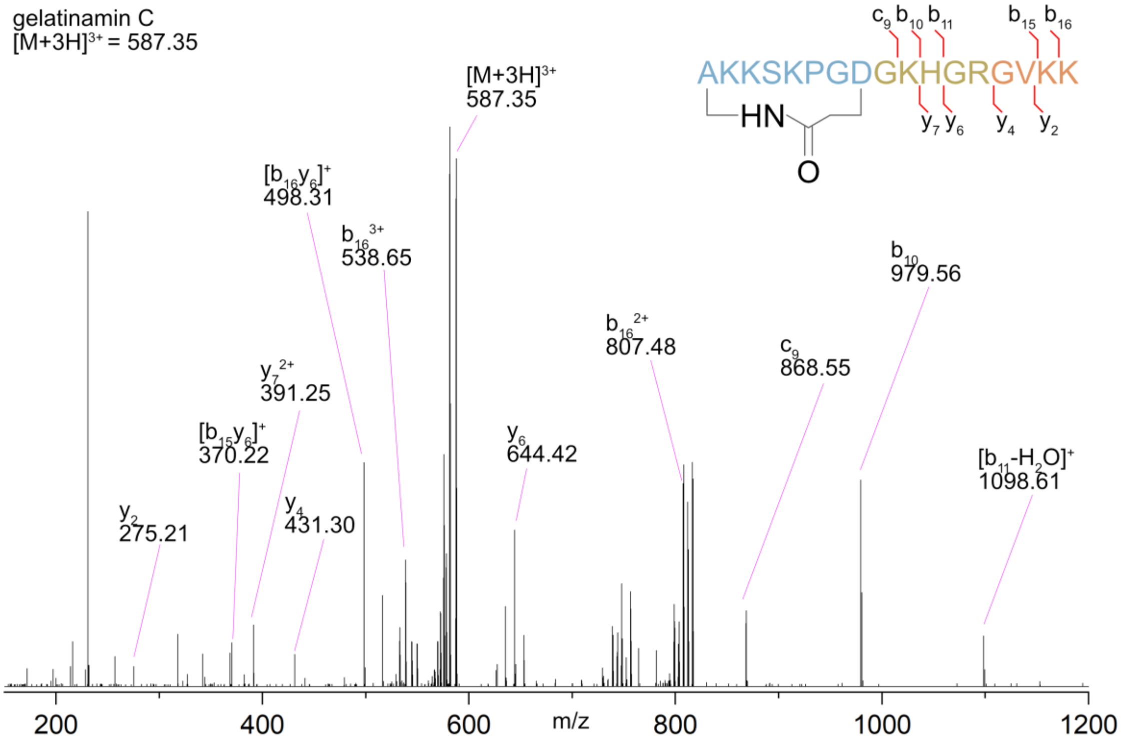


**Figure S14.** The MS/MS spectrum of gelatinamin C ($[M+3H]^{3+}$) obtained from the GelP reaction with gelatinamin A and glycine. The shown gelatinamin C sequence is not shown as a lasso peptide for simplicity.

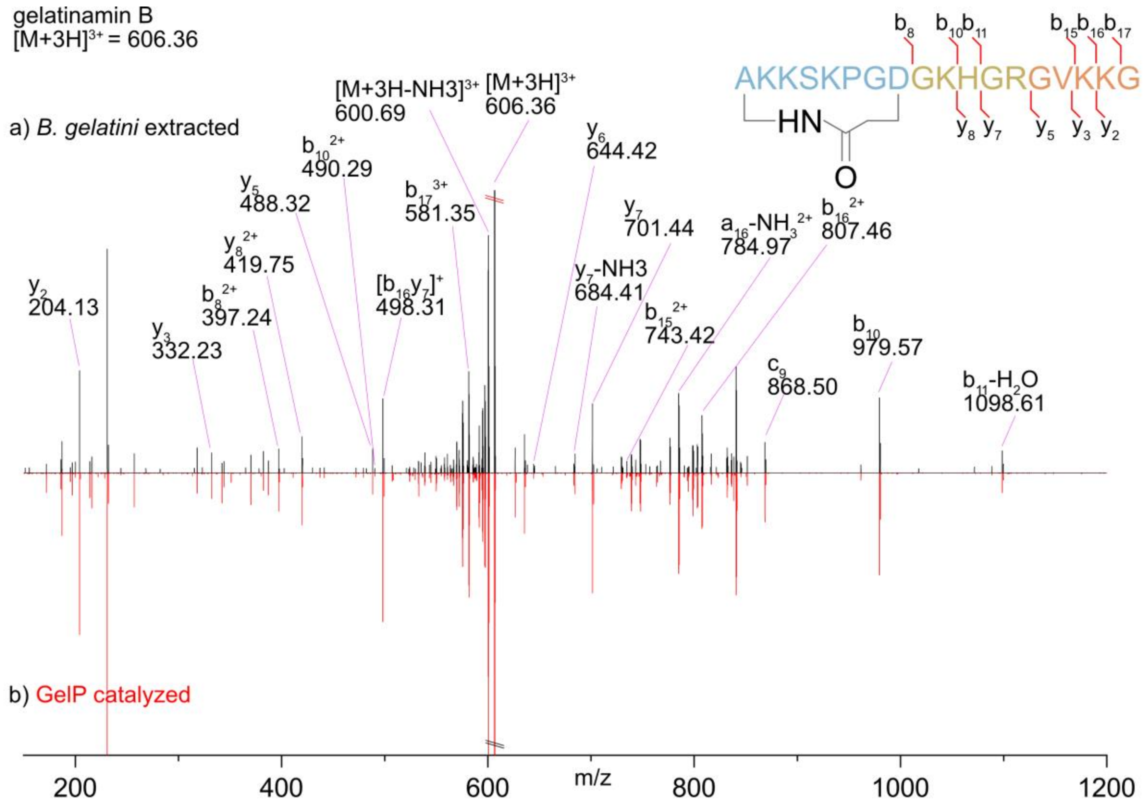


**Figure S15.** Comparison of MS/MS spectra of gelatinamin B ($[M+3H]^{3+}$]) obtained from the native producer *B. gelatini* and from GelP catalyzed reaction with gelatinamin A. The absence of N-terminal results in limited fragmentation with no b-fragments smaller than $b_8$. a) *B. gelatini* produced gelatinamin B MS/MS spectrum. b) GelP produced gelatinamin B MS/MS spectrum. The shown gelatinamin B sequence is not shown as a lasso peptide for simplicity.

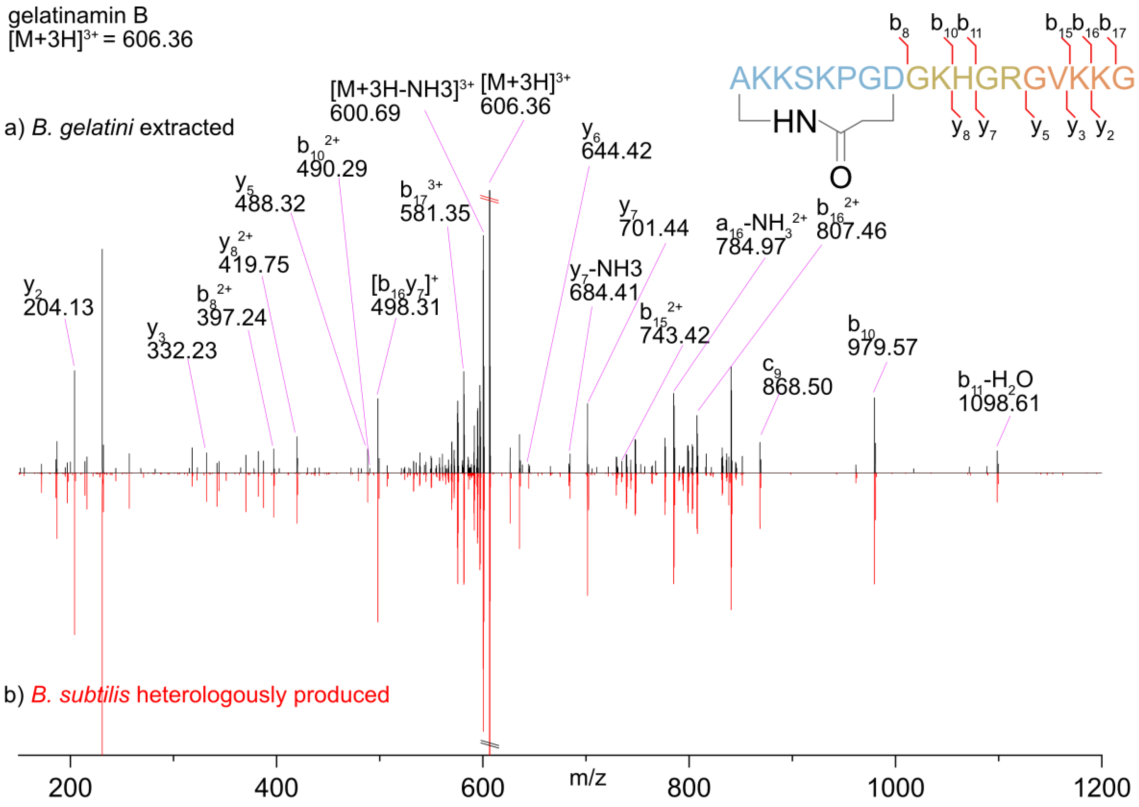


**Figure S16.** Comparison of MS/MS spectra of gelatinamin B ($[M+3H]^{3+}$]) obtained from the native producer *B. gelatini* and from heterologous host. The absence of N-terminal results in limited fragmentation with no b-fragments smaller than $b_8$. a) *B. gelatini* produced gelatinamin B MS/MS spectrum. b) *B. subtilis* produced gelatinamin B MS/MS spectrum. The shown gelatinamin B sequence is not shown as a lasso peptide for simplicity.

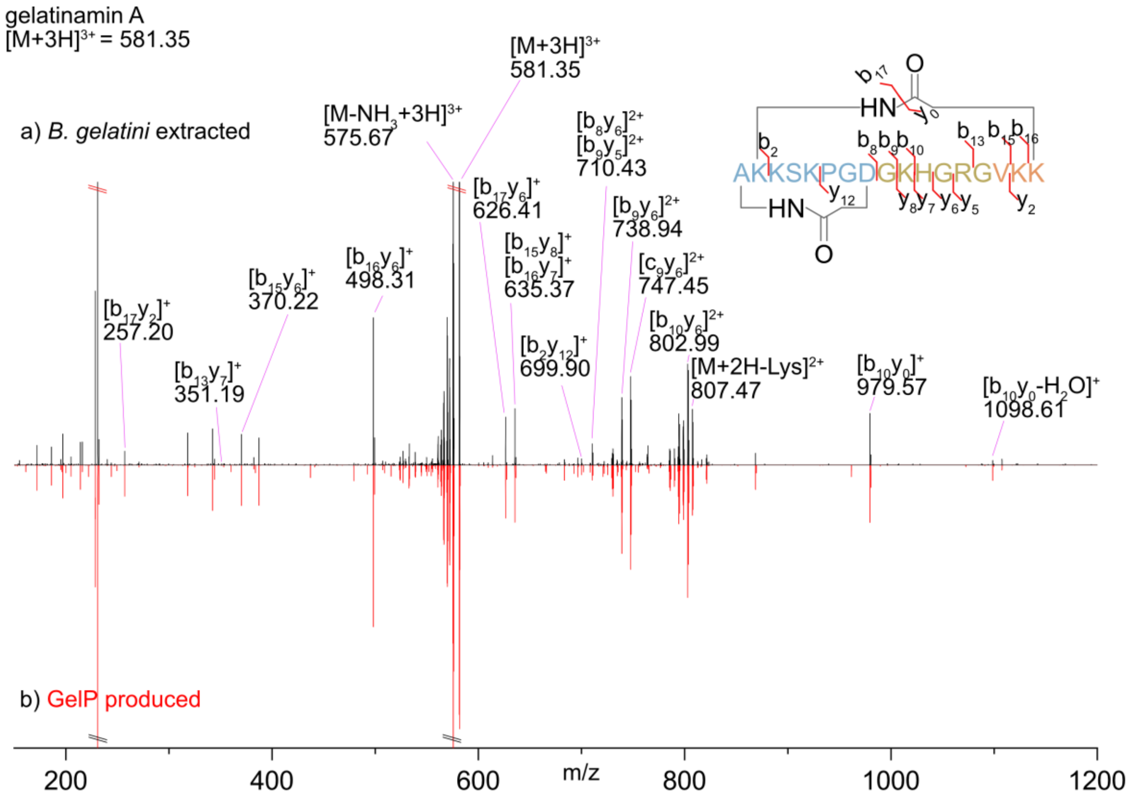


**Figure S17.** Comparison of MS/MS spectra of gelatinamin A ($[M+3H]^{3+}$]) obtained from the native producer *B. gelatini* and from GelP catalyzed reaction with gelatinamin B. The absence of both N- and C-terminus results in observation of only double fragments. a) *B. gelatini* produced gelatinamin A MS/MS spectrum. b) GelP produced gelatinamin A MS/MS spectrum. The shown gelatinamin A sequence is not shown as a lasso peptide for simplicity.

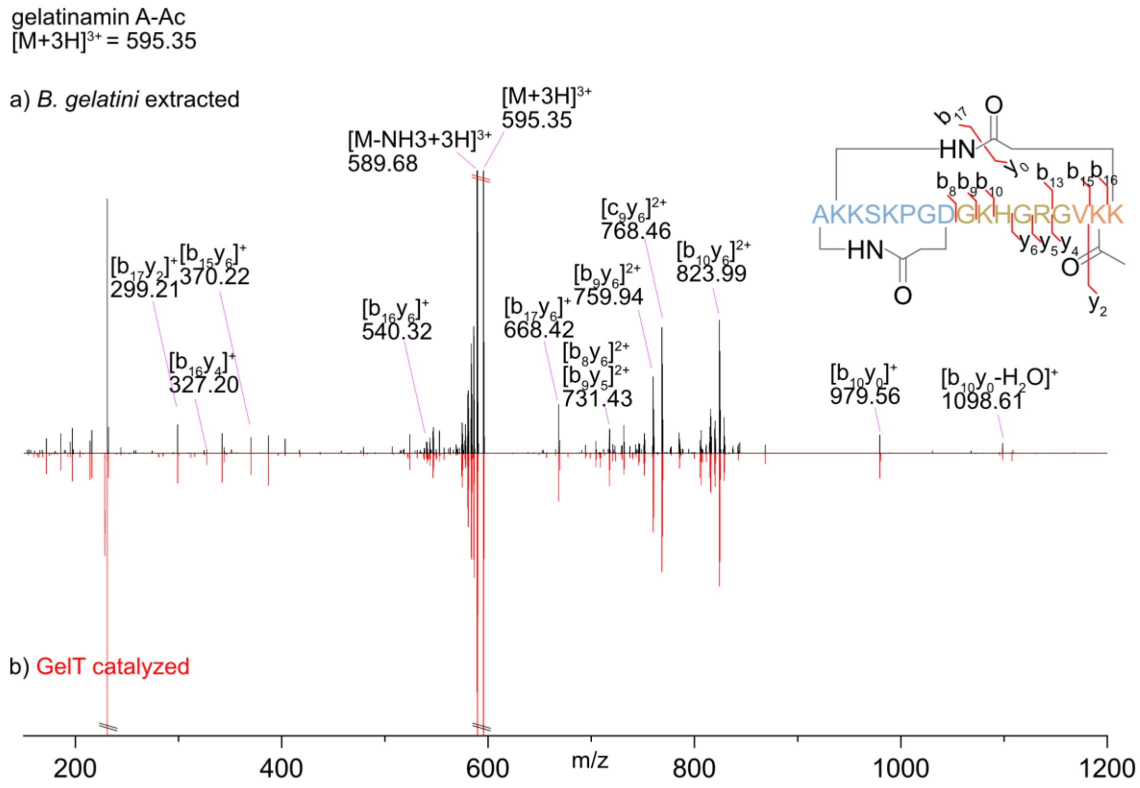


**Figure S18.** Comparison of MS/MS spectra of gelatinamin A-Ac ($[M+3H]^{3+}$) obtained from the native producer *B. gelatini* and from GelT catalyzed acetylation of gelatinamin A. a) *B. gelatini* produced gelatinamin A-Ac MS/MS spectrum. b) GelT acetylated gelatinamin A-Ac MS/MS spectrum. The shown gelatinamin A-Ac sequence is not shown as a lasso peptide for simplicity.

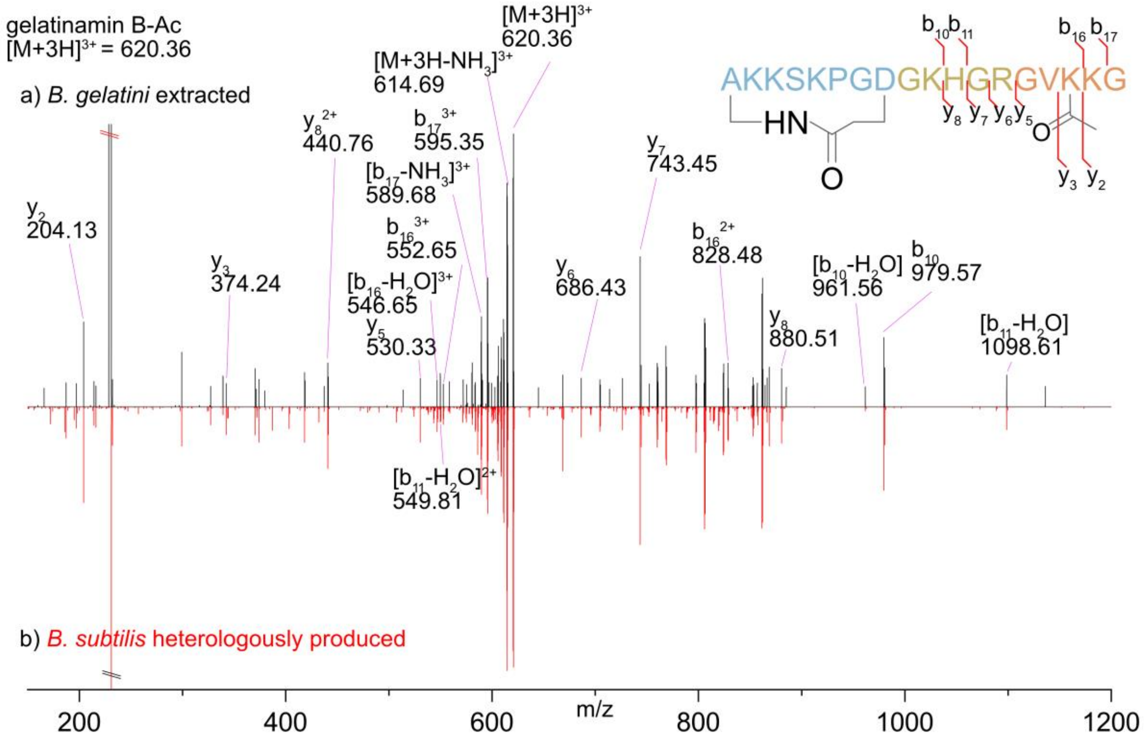


**Figure S19.** Comparison of MS/MS spectra of gelatinamin B-Ac ([M+3H]$^{3+}$]) obtained from the native producer *B. gelatini* and from heterologous host. a) *B. gelatini* produced gelatinamin B-Ac MS/MS spectrum. b) *B. subtilis* produced gelatinamin B-Ac MS/MS spectrum. The shown gelatinamin B-Ac sequence is not shown as a lasso peptide for simplicity.

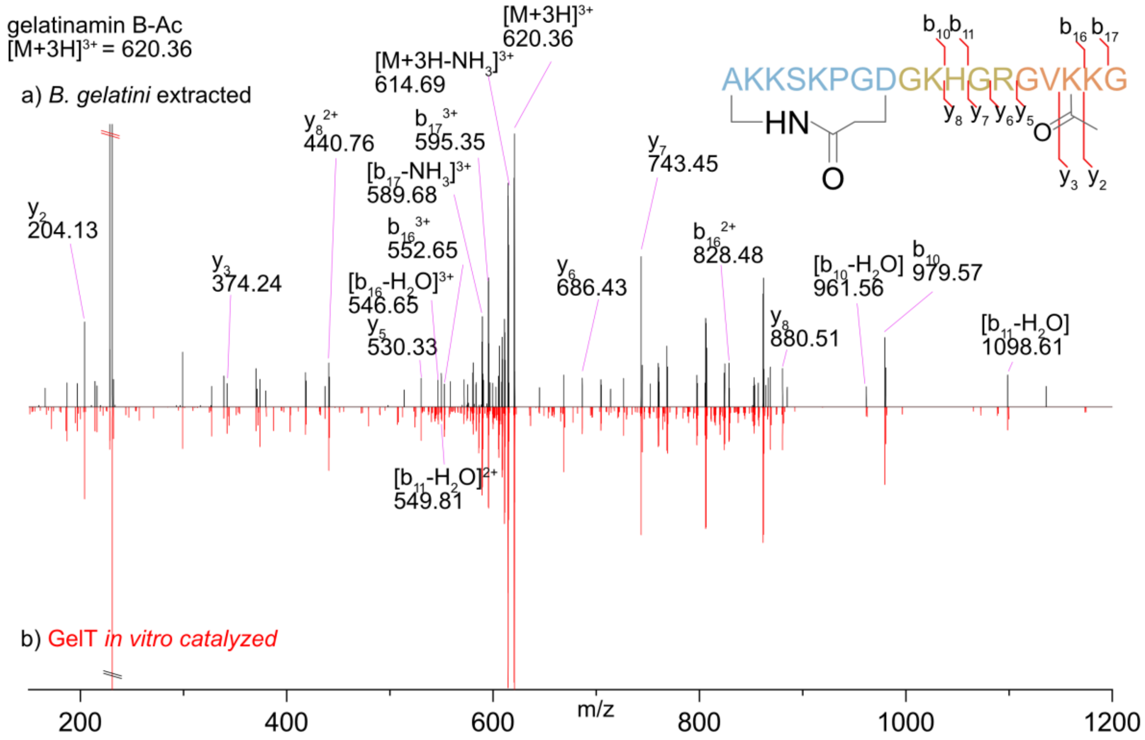


**Figure S20.** Comparison of MS/MS spectra of gelatinamin B-Ac ([M+3H]$^{3+}$]) obtained from the native producer *B. gelatini* and GelT catalyzed acetylation of gelatinamin B. a) *B. gelatini* produced gelatinamin B-Ac MS/MS spectrum. b) GelT acetylated gelatinamin B-Ac MS/MS spectrum. The shown gelatinamin B-Ac sequence is not shown as a lasso peptide for simplicity.

##### NMR supporting info

**Table S4.** List of recorded NMR spectra

| Experiment | Nuclei | Number of points (td) | Spectral Width f1 x f2 (ppm) | Scans | Mixing Time (ms) | Recycle Delay (s) |
|---|---|---|---|---|---|---|
| **TOCSY** | $^{1}H–^{1}H$ | 2048 x 1024 | 14 x 14 | 48 | 120 | 1.5 |
| **NOESY** | $^{1}H–^{1}H$ | 2048 x 1024 | 14 x 14 | 48 | 120 | 1.5 |
| **$^{15}N$ HSQC** | $^{1}H–^{15}N$ | 2048 x 256 | 14 x 34 | 64 | - | 1.5 |
| **$^{13}C$ HSQC** | $^{1}H–^{13}C$ | 2048 x 256 | 14 x 80 | 64 | - | 1.5 |

**Table S5.** NMR assignment Chemical shift assignment [ppm]. Asterisks (*) indicate ambiguity of geminal atoms and a double asterisks (**) indicate intraresidue ambiguity.

| AA | | NH | N | Hα | Hβ | Hγ | Hδ | Hε | Hζ | Cα | Cβ | Cγ | Cδ | Cε |
|---|---|---|---|---|---|---|---|---|---|---|---|---|---|---|
| 1 | Ala | 8.17 | 122.3 | 4.30 | 1.23 | - | - | - | - | 51.3 | 21.1 | - | - | - |
| 2 | Lys | 9.38 | 127.0 | 4.45 | 1.82, 1.74 | 1.32, 1.22, | 1.35, 1.17 | 3.44, 2.98 | 7.59* | 57.1 | 31.6 | 24.7 | 28.2 | 42.7 |
| 3 | Lys | 8.71 | 123.4 | 3.63 | 1.73, 1.65 | 1.36, 1.28 | 1.57, 1.52 | 2.91* | 7.49* | 57.2 | 35.4 | 25.4 | 28.6 | 42.2 |
| 4 | Ser | 8.20 | 112.0 | 3.88 | 3.98, 3.63 | - | - | - | - | 57.6 | 62.5 | - | - | - |
| 5 | Lys | 8.06 | 120.7 | 4.39 | 1.65, 1.53 | 1.22* | 1.53, 1.50 | 2.83* | 7.40* | 53.8 | 34.1 | 24.2 | 29.1 | 41.9 |
| 6 | Pro | - | - | 4.51 | 2.41, 2.21 | 1.85, 1.68 | 3.43, 3.35 | - | - | 61.9 | 35.3 | 24.8 | 50.8 | - |
| 7 | Gly | 8.52 | 105.7 | 4.59, 3.86 | - | - | - | - | - | 46.1 | - | - | - | - |
| 8 | Asp | 7.74 | 119.7 | 4.75 | 3.19, 1.95 | - | - | - | - | 50.6 | 40.0 | - | - | - |
| 9 | Gly | 8.49 | 109.3 | 4.26, 3.48 | - | - | - | - | - | 44.9 | - | - | - | - |
| 10 | Lys | 7.42 | 119.6 | 4.56 | 1.67, 1.65 | 1.32, 1.22 | 1.54* | 2.99, 2.83 | - | 53.8 | 33.1 | 24.7 | 28.9 | 42.2 |
| 11 | His | 8.95 | 123.3 | 4.22 | 3.17, 3.11 | - | 8.54** | 7.23** | - | 57.7 | 27.9 | - | - | - |
| 12 | Gly | 8.81 | 114.0 | 3.87, 3.57 | - | - | - | - | - | 45.6 | - | - | - | - |
| 13 | Arg | 7.74 | 119.3 | 4.54 | 1.92, 1.82 | 1.42, 1.35 | 3.06, 2.95 | 6.97 | - | 54.2 | 31.6 | 26.8 | 42.7 | - |
| 14 | Gly | 6.57 | 106.5 | 4.80, 3.52 | - | - | - | - | - | 45.8 | - | - | - | - |
| 15 | Val | 7.54 | - | 4.18 | 2.32 | 0.75*, 0.71* | - | - | - | 62.0 | 32.5 | 21.6, 18.0 | - | - |
| 16 | Lys | 8.38 | 121.2 | 4.34 | 1.63, 1.59 | 1.47, 1.31 | 1.57* | 3.00, 2.97 | 7.53* | 55.4 | 33.3 | 25.3 | 28.8 | 42.3 |
| 17 | Lys | 8.77 | 114.9 | 3.63 | 1.86, 1.80 | 1.29, 1.24 | 1.58, 1.52 | 2.88* | 7.47* | 57.3 | 29.4 | 25.3 | 29.1 | 42.1 |

**Table S6. List of Long-range NOE used by Cyana 2.1.**

| Residue | Long Range NOEs |
|---|---|
| Ala1 | #5 |
| Lys2 | #6 |
| Lys3 | #2 |
| Ser4 | #2 |
| Lys5 | #1 |
| Pro6 | #1 |
| Gly7 | #0 |
| Asp8 | #4 |
| Gly9 | #1 |
| Lys10 | #1 |
| His11 | #0 |
| Gly12 | #0 |
| Arg13 | #3 |
| Gly14 | #5 |
| Val15 | #5 |
| Lys16 | #3 |
| Lys17 | #1 |

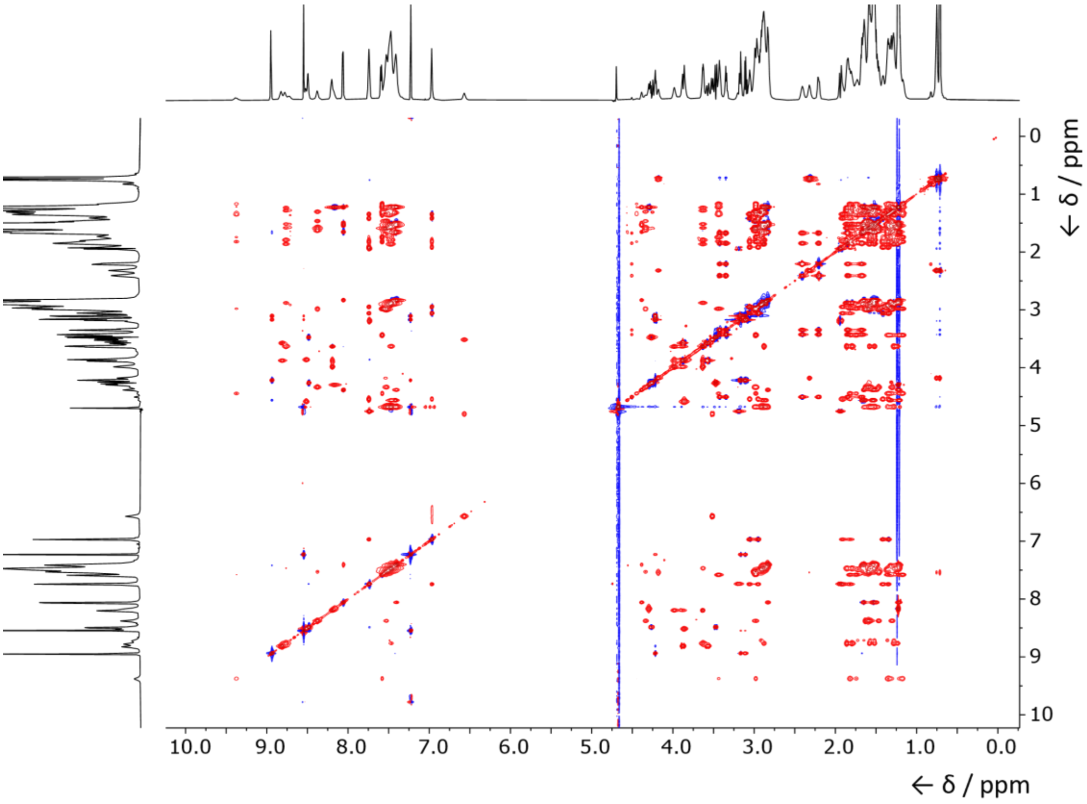


**Figure S21.** $^{1}$**H-**$^{1}$**H** TOCSY spectrum of gelatinamin A with 120 ms mixing time.

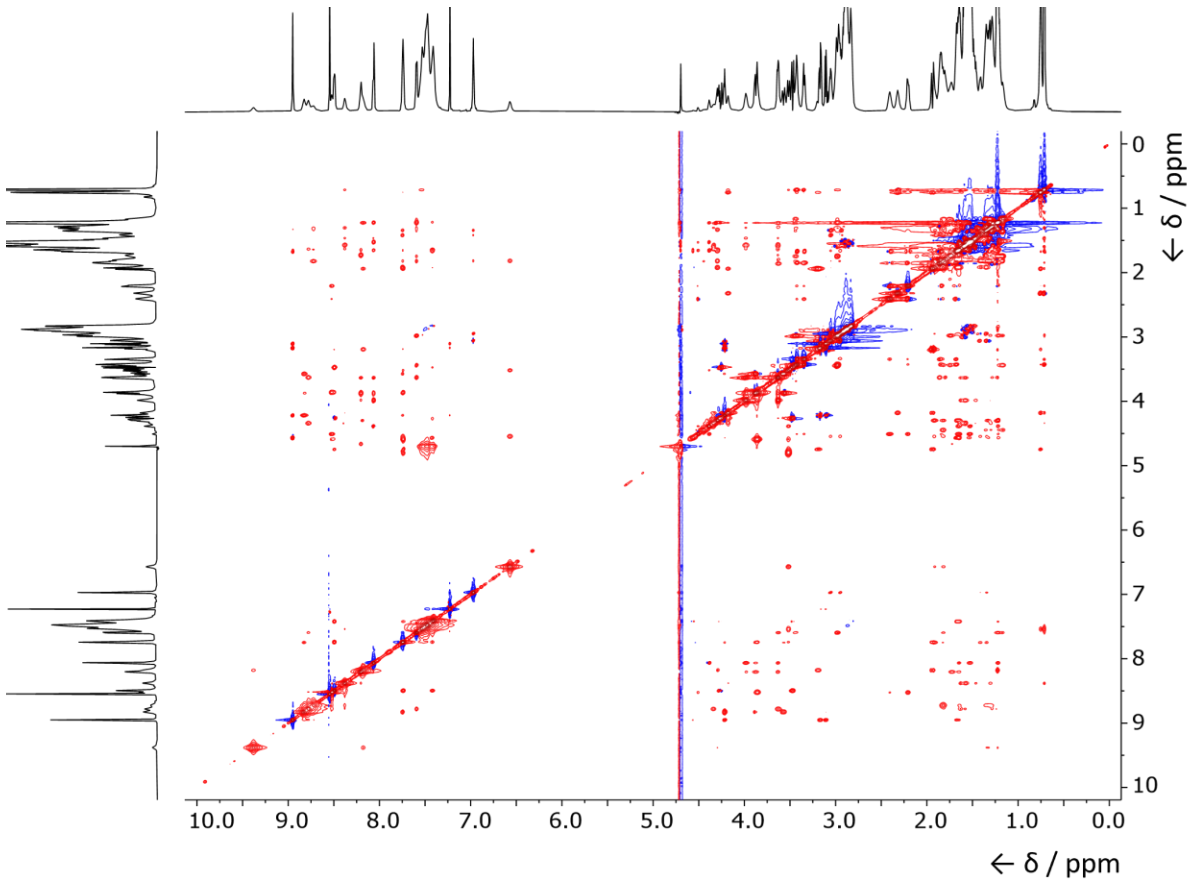


**Figure S22.** **[1]H-[1]H** NOESY spectrum of gelatinamin A with 120 ms mixing time.

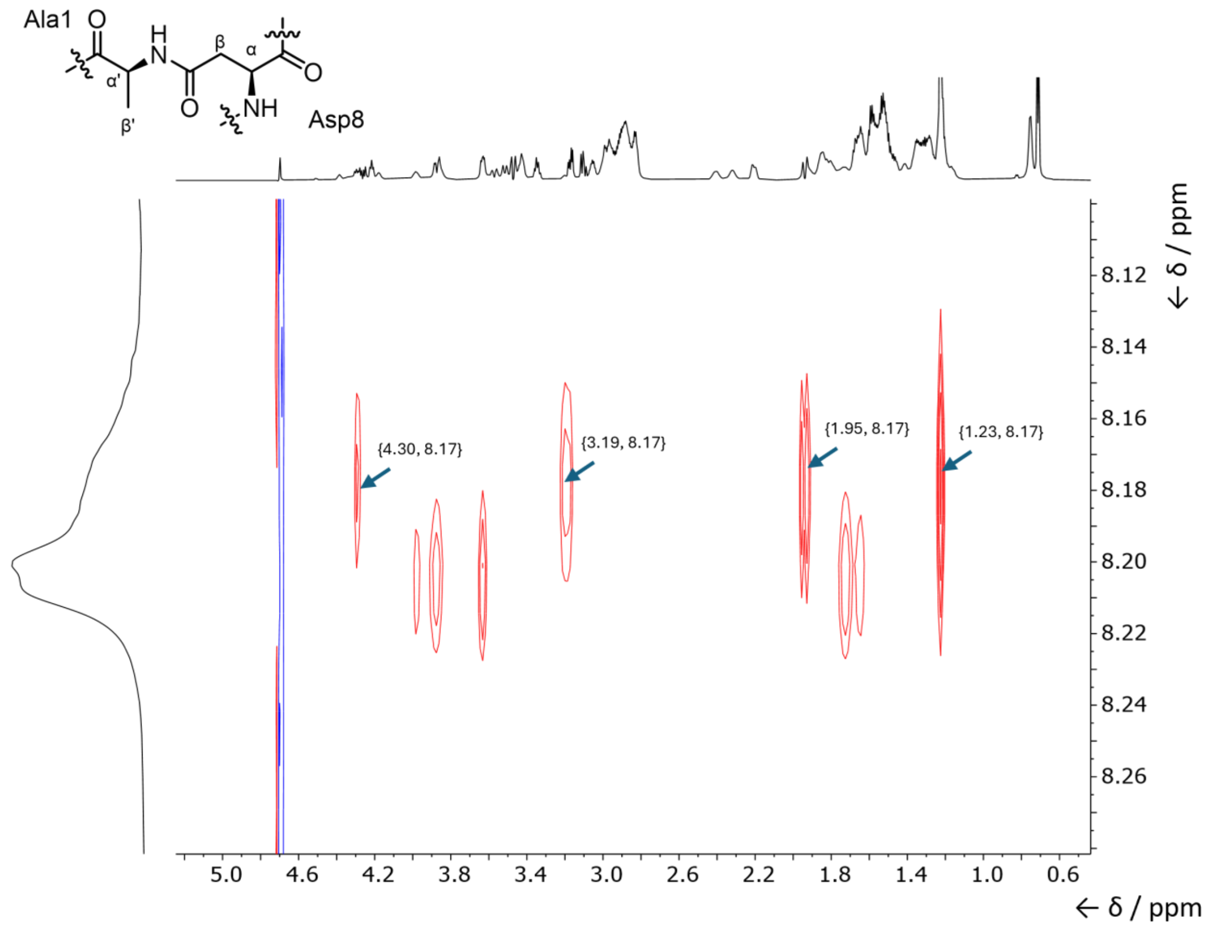


**Figure S23.** **$^1$H-$^1$H** NOESY (120 ms mixing time) correlations confirming the Ala1–Asp8 isopeptide linkage in gelatinamin A. Excerpt of the 120 ms NOESY spectrum showing the NOE cross-peaks from the Ala1 amide proton (δ 8.17 ppm) to aliphatic resonances assigned to Ala1 and Asp8. Correlations to Ala1 Hα (δ 4.30 ppm) and Hβ (δ 1.23 ppm) establish intra-residue proximity, while additional NOEs to Asp8 Hβ protons (δ 3.19 and 1.95 ppm), which appear as diastereotopic signals, demonstrate spatial contact between A1 NH and the D8 β-methylene group. Together, these NOE interactions support the presence of the A1–D8 macrolactam (isopeptide) bond characteristics of class V lasso peptides.

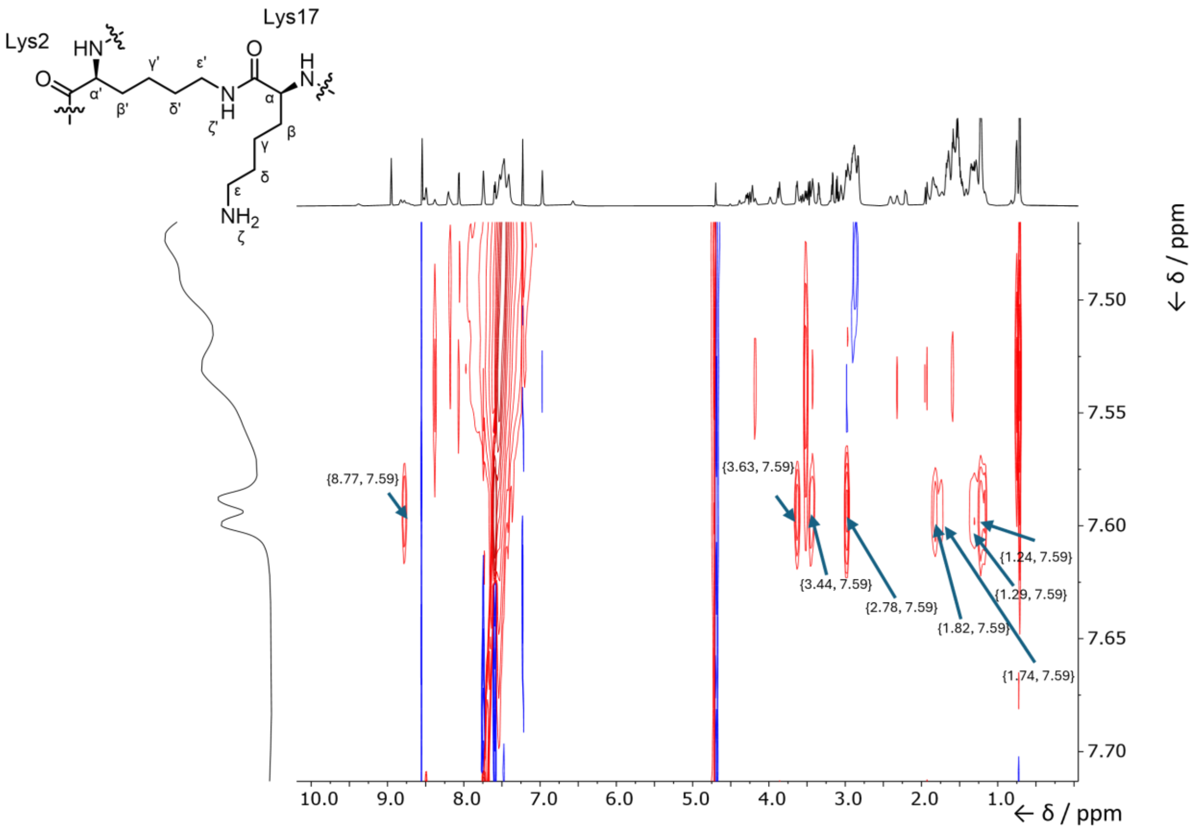


**Figure S24.** **$^1$H-$^1$H** NOESY (120 ms mixing time) correlations supporting the Lys2–Lys17 isopeptide linkage in gelatinamin A. Excerpt of the 120 ms NOESY spectrum showing the diagnostic NOE contacts of the isopeptide amide proton of Lys2 (δ 7.59 ppm), which is observed as a doublet and confirmed as part of the Lys2 spin system by TOCSY. This isopeptide NH exhibits a strong NOE to the backbone amide of Lys17 (δ 8.77 ppm), providing the most direct evidence for the Lys2–Lys17 macrolactam connection. Additional NOEs to Lys17 Hα (δ 3.63 ppm) and the Lys17 γ-methylene protons (δ 1.29 and 1.24 ppm) further support the spatial proximity of the two residues within the macrocycle. The isopeptide NH also shows correlations to its own side-chain resonances, including Lys2 Hε (δ 3.44 and 2.98 ppm) and Lys2 Hβ (δ 1.82 and 1.74 ppm), consistent with its assignment. Together, these NOE interactions confirm formation of the Lys2–Lys17 isopeptide bond.

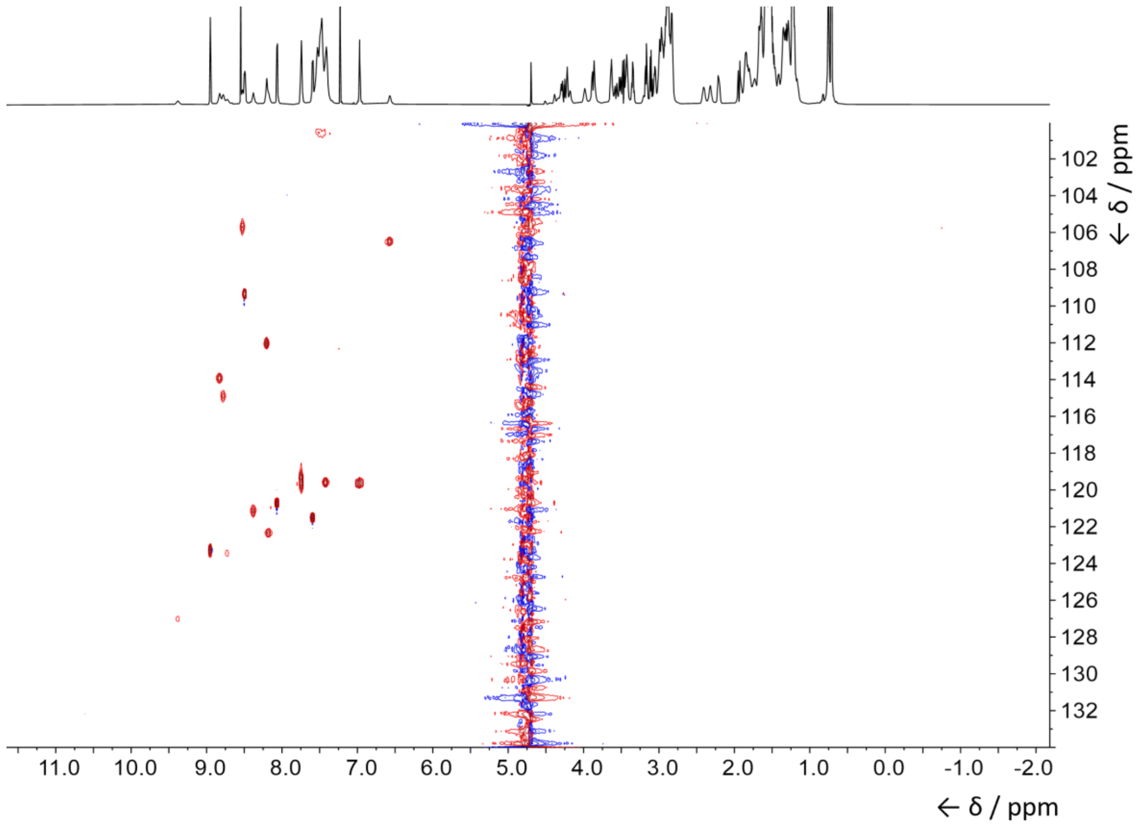


**Figure S25.** **[1]H-**[15]N HSQC spectrum of gelatinamin A.

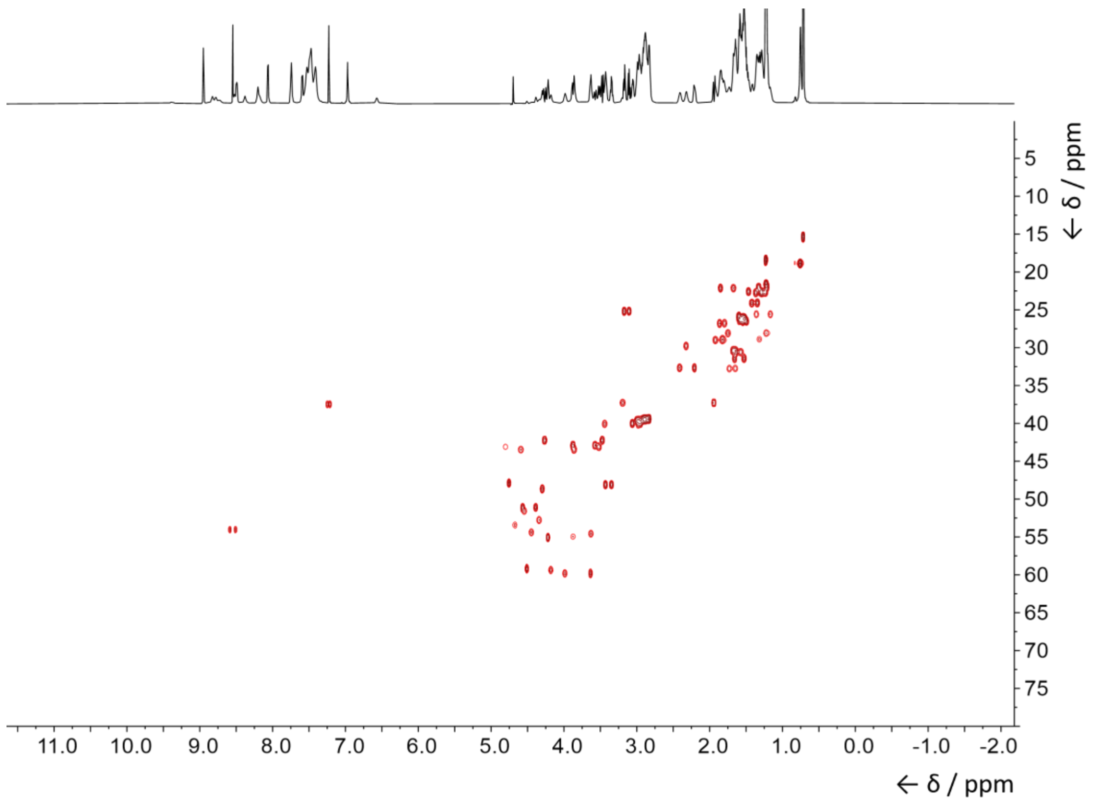


**Figure S26.** **[1]H-**[13]C HSQC spectrum of gelatinamin A.

[1] W. F. Vranken, W. Boucher, T. J. Stevens, R. H. Fogh, A. Pajon, M. Llinas, E. L. Ulrich, J. L. Markley, J. Ionides, E. D. Laue, “The CCPN data model for NMR spectroscopy: Development of a software pipeline” *Proteins: Structure, Function, and Bioinformatics* **2005**, *59*, 687–696.

[2] A. Merrild, T. Svenningsen, M. G. Chevrette, T. Tørring, “Evolution-Guided Discovery of Antimycobacterial Triculamin-Like Lasso Peptides” *Angewandte Chemie International Edition* **2025**, *64*, e202425134.

[3] X. Zhang, Y. -H. P. Zhang, “Simple, fast and high-efficiency transformation system for directed evolution of cellulase in *Bacillus subtilis*” *Microbial Biotechnology* **2011**, *4*, 98–105.

[4] F. Teufel, J. J. Almagro Armenteros, A. R. Johansen, M. H. Gíslason, S. I. Pihl, K. D. Tsirigos, O. Winther, S. Brunak, G. von Heijne, H. Nielsen, “SignalP 6.0 predicts all five types of signal peptides using protein language models” *Nat Biotechnol* **2022**, *40*, 1023–1025.

[5] J. Abramson, J. Adler, J. Dunger, R. Evans, T. Green, A. Pritzel, O. Ronneberger, L. Willmore, A. J. Ballard, J. Bambrick, S. W. Bodenstein, D. A. Evans, C.-C. Hung, M. O’Neill, D. Reiman, K. Tunyasuvunakool, Z. Wu, A. Žemgulytė, E. Arvaniti, C. Beattie, O. Bertolli, A. Bridgland, A. Cherepanov, M. Congreve, A. I. Cowen-Rivers, A. Cowie, M. Figurnov, F. B. Fuchs, H. Gladman, R. Jain, Y. A. Khan, C. M. R. Low, K. Perlin, A. Potapenko, P. Savy, S. Singh, A. Stecula, A. Thillaisundaram, C. Tong, S. Yakneen, E. D. Zhong, M. Zielinski, A. Žídek, V. Bapst, P. Kohli, M. Jaderberg, D. Hassabis, J. M. Jumper, “Accurate structure prediction of biomolecular interactions with AlphaFold 3” *Nature* **2024**, *630*, 493–500.

[6] H. Nielsen in *Large Language Models (LLMs) in Protein Bioinformatics* (Ed.: D.B. Kc), Springer US, New York, NY, **2025**, pp. 153–175.

[7] P. Natale, T. Brüser, A. J. M. Driessen, “Sec- and Tat-mediated protein secretion across the bacterial cytoplasmic membrane—Distinct translocases and mechanisms” *Biochimica et Biophysica Acta (BBA) - Biomembranes* **2008**, *1778*, 1735–1756.

[8] K. Jeanne Dit Fouque, H. Lavanant, S. Zirah, J. D. Hegemann, C. D. Fage, M. A. Marahiel, S. Rebuffat, C. Afonso, “General rules of fragmentation evidencing lasso structures in CID and ETD” *Analyst* **2018**, *143*, 1157–1170.
